# In Vivo Transcriptomics-Based Framework for Systematic Indication Expansion

**DOI:** 10.64898/2026.10.12.758524

**Authors:** Zsolt Palmer, Shubham Koirala, Nabasmita Talukdar, Xiaodan Zhang, Hannah Garver, McKenzie Mahlmeister, Li Huang, Dimitri Joseph, Dmitry Leshchiner, Rama Shankar, Stephanie W. Watts, Xiaopeng Li, Benjamin S. Glicksberg, Adam Lauver, Bin Chen

**Affiliations:** Department of Biochemistry and Molecular Biology, College of Natural Science, Michigan State University; East Lansing, MI, USA; Genetics and Genome Sciences Program, Michigan State University; East Lansing, MI, USA; Department of Epidemiology and Biostatistics, Michigan State University; East Lansing, MI, USA; Department of Pediatrics and Human Development, Michigan State University; East Lansing, MI, USA; Department of Pharmacology and Toxicology, College of Human Medicine, Michigan State University; Grand Rapids, MI, USA; Icahn School of Medicine at Mount Sinai; New York, NY, USA; Center for AI-enabled Drug Discovery, Michigan State University; Grand Rapids, MI 49503, USA

## Abstract

Capturing the systemic effects of drugs *in vivo* remains a fundamental challenge for drug repurposing strategies. Here, we present an *in vivo* transcriptomics-based drug repurposing framework that profiles multi-organ RNA-seq from drug-treated rodents to model systemic drug responses and match tissue-specific drug signatures against a compendium of disease signatures for indication expansion. We applied this framework to semaglutide, a GLP-1 receptor agonist originally developed for diabetes and subsequently expanded for several other indications. By harmonizing more than 900 human disease transcriptomic signatures via agentic AI and using Z-transformed reversal of gene expression scoring to quantify drug-disease relationships, our approach successfully recovered established semaglutide indications while deprioritizing Alzheimer’s disease, an indication for which semaglutide was not effective. Top predictions for common diseases were supported by real-world electronic health record data from the Mount Sinai Health System and several predictions for rare indications are proposed for further investigation. This framework provides a scalable strategy for indication expansion across diverse therapeutic agents, especially investigational compounds for which real-world evidence remains sparse.

**One Sentence Summary:** Comparing drug and disease expression profiles from *in vivo* contexts achieves indication profiles reflective of the interconnected mammalian organism.

## INTRODUCTION

Indication expansion, a form of drug repurposing, is becoming an increasingly common strategy in pharmaceutical development (*1–3*). In oncology, more than 50% of approved drugs hold approval for more than one indication (*4, 5*), but the trend extends beyond just oncology, driven by repurposing’s cost-effectiveness, reduced development timelines, and ability to fill commercial gaps for the pharmaceutical industry. Transcriptomic reversal is a widely adopted computational repurposing/expansion approach, particularly for first-pass screening (*6, 7*). The central principle, established by the Connectivity Map (*8*) and extended by the LINCS-L1000 initiative (*9*) and others (*10–18*), holds that a drug capable of reversing the transcriptomic signature of a disease represents a viable therapeutic candidate. Transcriptomic reversal carries a particular advantage for rare diseases and newly approved or investigational drugs, for which electronic health record (EHR)-based approaches are often underpowered due to limited patients and sparse real-world evidence (*19*). This is especially pertinent given that thousands of rare diseases collectively affect over 260 million people worldwide, yet effective therapies remain unavailable for the majority (*20*). Rare diseases also represent an attractive opportunity for drug repurposing: orphan drug designation reduces regulatory burden (*21*), and approved drugs face a comparatively shorter path to new indications than novel compounds. Against a backdrop where 90% of drugs entering clinical development fail, the economic rationale for drug repurposing becomes especially compelling (*22*). Applying transcriptomic reversal to this end has, however, been constrained by scale, quality and contextual relevance of available drug- and disease-associated transcriptomic signatures.

Existing drug-induced transcriptomic profiles are primarily derived in homogeneous, immortalized cell lines (*8, 9*), which fail to capture the pleiotropic effects of drugs across the heterogeneous cellular environments of tissues and the coordinated physiology of the mammalian organism. This limitation is particularly consequential for drugs with notable pleiotropic mechanisms such as glucagon-like peptide-1 receptor agonists (GLP-1 RAs), whose effects span across tissues through both direct receptor engagement and indirect systemic mechanisms, resulting in widespread transcriptional and physiological responses (*23, 24*).

Previous transcriptomic repurposing efforts have focused primarily on identifying treatments for individual diseases (*13, 17, 18*). Efficient drug-centric indication expansion requires a systematic and unbiased assessment of a drug’s therapeutic potential across diverse diseases. Such analyses have been limited by the scale and contextual relevance of available disease-associated transcriptomic signatures. To obtain large-scale *in vivo* signatures, many turn to transcriptomic repositories such as the Gene Expression Omnibus (GEO) (*25*) which now indexes millions of samples across hundreds of thousands of studies, and is uniformly quantified in resources such as ARCHS4 (*26*) and recount3 (*27*), providing extraordinary breadth across diseases, tissues, and experimental designs (*28*). Some existing disease-signature resources demonstrate the value of these transcriptomic atlases; RummaGEO (*29*) computed differential expression signatures through automatically identified groups of samples that correspond to conditions in GEO studies, and DiSignAtlas provides a broad collection of human and mouse disease signatures from bulk and single-cell transcriptomic studies (*30*), but these repositories highlight certain gaps for applicability to transcriptomic reversal. Disease signatures for drug-centric repurposing require robust processing, standardized disease and control labeling, tissue matching, and exclusion of treated or non-primary samples. Many public disease studies lack matched healthy controls, use controls that differ in tissue source or processing context, or mix treated and untreated samples in ways that confound disease-state modeling. Moreover, for systemic reversal screening, the requirements are yet more stringent: signatures must be traceable to disease-relevant tissues, filtered for sample context, and generated in a way that supports tissue-aware pairing with drug perturbation profiles.

Here, we present a framework for an unbiased, programmatic comparison of the *in vivo* transcriptomic states induced by drugs and diseases across tissues (Fig. 1A). First, we leveraged the GTEx reference RNA-seq database, ARCHS4, a large collection of processed GEO RNA-seq samples, and GEOMeta (*31*), a GEO sample metadata resource generated using agentic AI, to construct a harmonized human *in vivo* disease transcriptomic reference atlas comprising 979 disease signatures across 353 diseases and 48 tissues. Second, rather than rely on existing perturbational profiles, which were derived overwhelmingly from cell lines and did not capture cross-tissue effects, we generated dedicated multi-organ transcriptomic profiles of semaglutide designed specifically to resolve its systemic action. Primary RNA-seq was collected from five organs (liver, heart, lung, kidney, and brain) of high-fat diet rats, and augmented with complementary mouse datasets from GEO, yielding 35 *in vivo* signatures spanning the tissues where semaglutide acts (fig. S1A). By computing Z-transformed Reversal of Gene Expression Scores (RGES) between tissue-matched drug and disease signatures, we screened semaglutide against 108 diseases, including rare conditions for which no EHR-based screening approach would be feasible, producing a defined drug-disease landscape which nominates several novel repurposing candidates of interest.

**Fig. 1.**
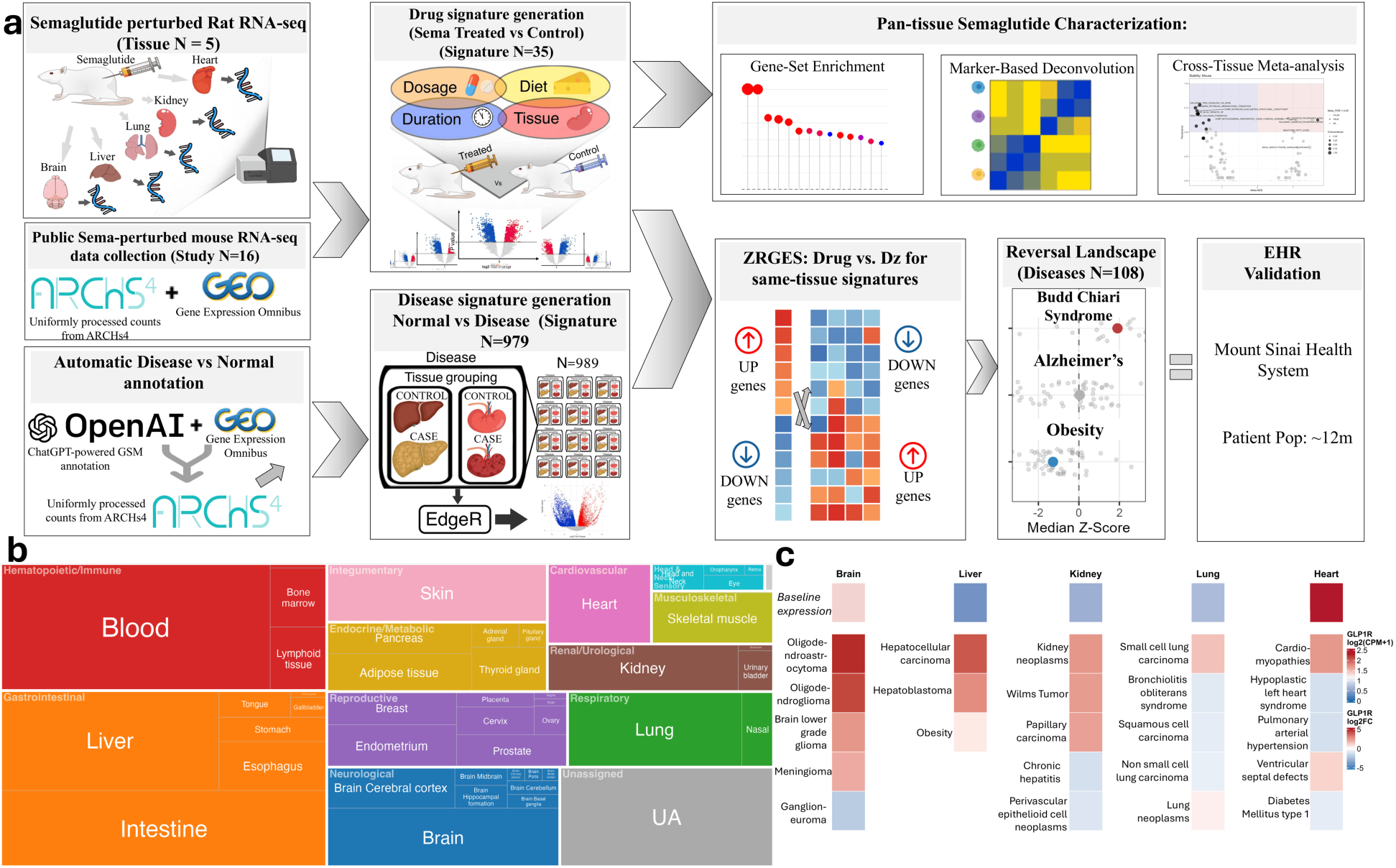
Overview of the in vivo transcriptomics-based indication expansion framework. **(A)** Schematic overview of the transcriptomic indication expansion framework. **(B)** Treemap displaying the tissue source distribution of all disease signatures in the atlas, grouped by organ system. Tile area is proportional to the number of signatures derived from each tissue, and tiles are colored by organ system category. The atlas spans 48 distinct tissue types across 13 organ systems, with the largest representation from hematopoietic/immune, gastrointestinal, and neurological tissues, reflecting the breadth of publicly available disease transcriptomic data. **(C)** GLP1R baseline expression and disease-associated dysregulation across five example tissues. Each column represents one tissue (brain, liver, kidney, lung, heart). The top box (“Baseline expression”) shows mean GLP1R transcript abundance in normal control tissue, computed as log2(CPM+1) from control samples across GEO series in ARCHS4, on an absolute blue-to-red scale (blue = low, red = high). Below, each column shows the five diseases with the largest absolute GLP1R log2 fold change in that tissue, ranked top to bottom by effect sizes, colored on a separate diverging scale (blue = down-regulated, red = up-regulated relative to matched controls; values capped at ±5). Note that the two scales represent different quantities, absolute baseline expression versus relative disease-associated change, and are not directly comparable. Such analysis can be carried out for any receptor gene, not merely GLP1R. Tissues were selected to enable analysis in relation to semaglutide signatures. Metastatic and circulating RNA disease signatures were removed from this graph for legibility.

## RESULTS

### *In vivo* drug- and disease-associated transcriptomic signatures

A compendium of high-quality human disease signatures was generated using sources from GEO (*25*), GTEx (*32*), and Treehouse (*33, 34*), containing 979 unique signatures across 353 diseases. The compendium spans a broad landscape of human pathology, encompassing signatures derived from 48 distinct tissue types organized across 13 organ systems, with the largest representation from hematopoietic/immune, gastrointestinal, and neurological tissues, reflecting the breadth of publicly available disease transcriptomic data (Fig. 1B). To assess disease signature quality, we evaluated their concordance with DiSignAtlas, an independent disease signature dataset derived from GEO (fig. S1B). Because DiSignAtlas reports only a truncated set of differentially expressed (DE) genes for each signature (10-1,000 genes), the overlapped genes for correlation analysis are limited. Despite this constraint, disease signatures showed concordance, with a median Spearman correlation of 0.54 and 53.2% of signatures achieving r ≥ 0.5. Spearman correlation of signatures revealed consistent clustering by disease type (fig. S1C), and further, concordance between enriched pathways and generally known disease mechanisms were manually evaluated for disease groupings such as cancers, or respiratory diseases. The most enriched pathways for cancers, for instance, were cell cycle/proliferation pathways (fig. S1D, table S1).

To illustrate the biological insights enabled by the resource, we examined GLP1R expression across normal tissues and its dysregulation across diseases relative to matched controls (Fig. 1C), excluding metastases and circulating RNA for clarity. As expected, GLP1R had relatively high expression in the brain (*35*), whereas expression was relatively low in the liver, kidney and lung. Interestingly, expression in the heart was the highest amongst these organs, despite a contentious expression profile in this tissue (*36, 37*). Semaglutide’s relationship to a variety of cancers has been an area of some focus (*38*), and GLP1R was significantly dysregulated across a broad range of cancers. For instance, it was upregulated in low-grade glioma, where higher expression has been associated with improved overall survival (*39*). We also identified marked GLP1R dysregulation in additional disease contexts, including oligodendroastrocytoma, highlighting opportunities to investigate their implications in these disease contexts.

With the disease signatures prepared, we next captured semaglutide’s systemic action across major organ systems by generating bulk RNA-seq in a high-fat diet rat model and supplemented these with curated mouse datasets, yielding 35 *in vivo* signatures for downstream analyses (table S2). Pathway enrichment analyses across the Hallmark, Reactome, and Gene Ontology: Biological Processes (GO:BP) libraries revealed a conserved anti-inflammatory and extracellular-matrix suppressive transcriptional core across tissues (Fig. 2A, B, fig. S2A, B, table S3). Meta-analysis of these semaglutide-perturbed signatures (signed Stouffer combination across signatures, per database) identified a set of pathways concordantly enriched in ≥ 90% of signatures and significant after Benjamini-Hochberg correction (tables S4, S5). This conserved core was dominated by the downregulation of immune and matrix-remodeling programs such as the interferon gamma response (Hallmark, FDR = 1 × 10^−112^), inflammatory response (Hallmark, FDR = 2.4 × 10^−61^), extracellular matrix organization (Reactome, FDR = 5 × 10^−147^), collagen fibril assembly (Reactome FDR = 2.55 × 10^−112^), and leukocyte migration, chemotaxis, and cytokine production (GO:BP, FDR < 0.05). Leave-one-out analysis confirmed the core was not driven by any individual signature: all five Hallmark, 26 of 28 Reactome, and 154 of 156 GO:BP core pathways remained significant across every iteration.

**Fig. 2.**
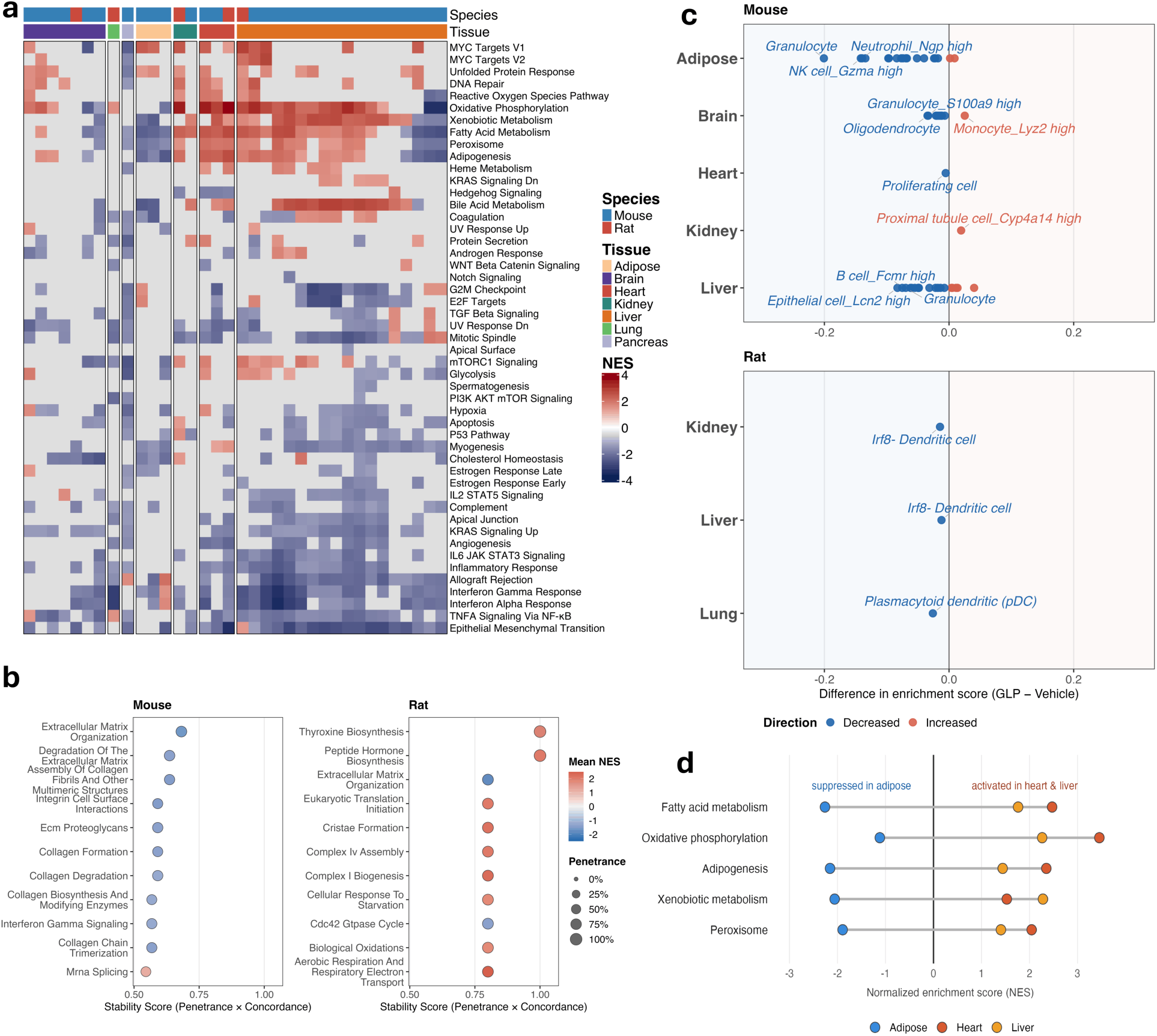
Semaglutide signature characteristics across tissues. **(A)** Heatmap of Hallmark gene set enrichment scores across the 35 semaglutide-perturbed transcriptomic signatures. Rows represent Hallmark pathways and columns represent individual signatures, both ordered by hierarchical clustering within tissue. Color indicates NES, with red denoting pathway activation (positive NES) and blue denoting suppression (negative NES). Column annotations indicate species and tissue of origin. **(B)** Pathway stability analysis of the most reproducibly enriched pathways across semaglutide-perturbed signatures in mouse and rat contexts, from the Reactome database. Pathways are ranked by a stability score defined as the product of prevalence (significantly enriched) and directional concordance of NES across signatures. Dot color indicates mean NES (red, activation; blue, suppression) and dot size indicates prevalence; the number of signatures significantly enriched in the same direction. **(C)** Marker-based deconvolution results showing estimated differences in cell type enrichment scores between semaglutide-treated and vehicle-treated animals across tissues in mice and rats. Dot color indicates direction of change (red, increased; blue, decreased), and labeled points represent the three cell types per tissue with the greatest mean effect across signatures by Fisher’s combined probability method. Raw p < 0.05. **(D)** NES values for hallmark pathways between (heart and liver) and white adipose tissue, Leave-One-Out tested for robustness.

Given that most signatures were derived from rodents on high-fat diets, part of the observed immune suppression possibly reflects reduced adipose inflammatory burden. To confirm this, we performed marker-based deconvolution of the bulk RNA-seq signatures and observed consistent downregulation of genes associated with immune cells. In rats, dendritic cell markers were broadly reduced across tissues, indicating suppression of a key cellular bridge between the innate and adaptive immune systems (Fig. 2C). In mice, a similar pattern emerged, including the downregulation of dendritic cells, granulocytes, and macrophages. To determine if this effect was specific to semaglutide, we compared the signatures between semaglutide and sodium/glucose cotransporter-2 inhibitors (SGLT2 inhibitors), a drug class approved for type 2 diabetes. Both drug classes affect the same immune cell populations similarly without significant differences in population changes (fig. S2C). This conserved and uniformly immune suppressive nature cannot be fully explained by weight loss alone, which incompletely resolves the adipose immune phenotype even after substantial fat loss (*40*). This suggests a direct or systemic immunomodulatory component of their actions.

In contrast to the conserved pathways, semaglutide also induced pronounced tissue-specific responses, primarily driven by opposing metabolic programs in white adipose tissue relative to other tissues (table S6). The most robust discordance was between white adipose tissue and both liver and heart tissues (Fig. 2D). In particular, lipid- and oxidative-metabolism programs were significantly upregulated in heart and liver but downregulated in white adipose tissue, with each cohort independently significant: fatty-acid metabolism (heart +2.47 Normalized Enrichment Score (NES) vs adipose −2.26 NES; both FDR < 10⁻^20^), and adipogenesis (+2.35 NES vs −2.16 NES; both FDR < 10^−20^).

### Semaglutide indication landscape characterized by *in vivo* transcriptomic reversal

We next characterized the semaglutide indication landscape by matching the drug- and disease-associated signatures by tissue, and calculating Z-transformed RGES. The raw RGES was converted to a Z-score against an empirical null of RGES values from randomly sampled, size-matched query gene sets. The Z-score expresses the strength of reversal relative to chance (Fig. 3, table S7). To highlight rare diseases, we estimated associated disease population sizes using data from the Mount Sinai Health System (fig. S3). To compute significance, Stouffer’s method was used to determine if a drug-induced signature significantly reversed or induced a disease signature, with non-significant results indicating a lack of directional effect. The framework successfully identified significant reversal effects for most approved semaglutide indications for which disease signatures were available, including metabolic dysfunction-associated steatohepatitis (MASH)(Z-RGES: -0.672, FDR = 1.08 × 10^−7^), obesity (Z-RGES: -1.174, FDR = 1.65 × 10^−12^), type 2 diabetes mellitus (Z-RGES: -1.854, FDR = 0.034), and heart failure (Z-RGES: -0.388, FDR = 8.1 × 10^−7^), indicating strong positive predictive power. Moreover, the framework did not predict significant reversal for diseases in which GLP-1 receptor agonists have not demonstrated established clinical benefit, including Alzheimer’s (Z-RGES: 0.042, FDR = 0.07), Parkinson’s (Z-RGES: 0.095, FDR = 0.9), and major depressive disorder (Z-RGES: 0.389, FDR = 0.02) (*41–43*). As a positive control to demonstrate that neurodegenerative disease signatures can be reversed, we collected GEO transcriptomic profiles for memantine, an FDA-approved Alzheimer’s treatment. Memantine showed significant aggregate reversal of the Alzheimer’s disease signature (median Z-RGES -0.371, FDR = 0.0064) (fig. S4A).

**Fig. 3.**
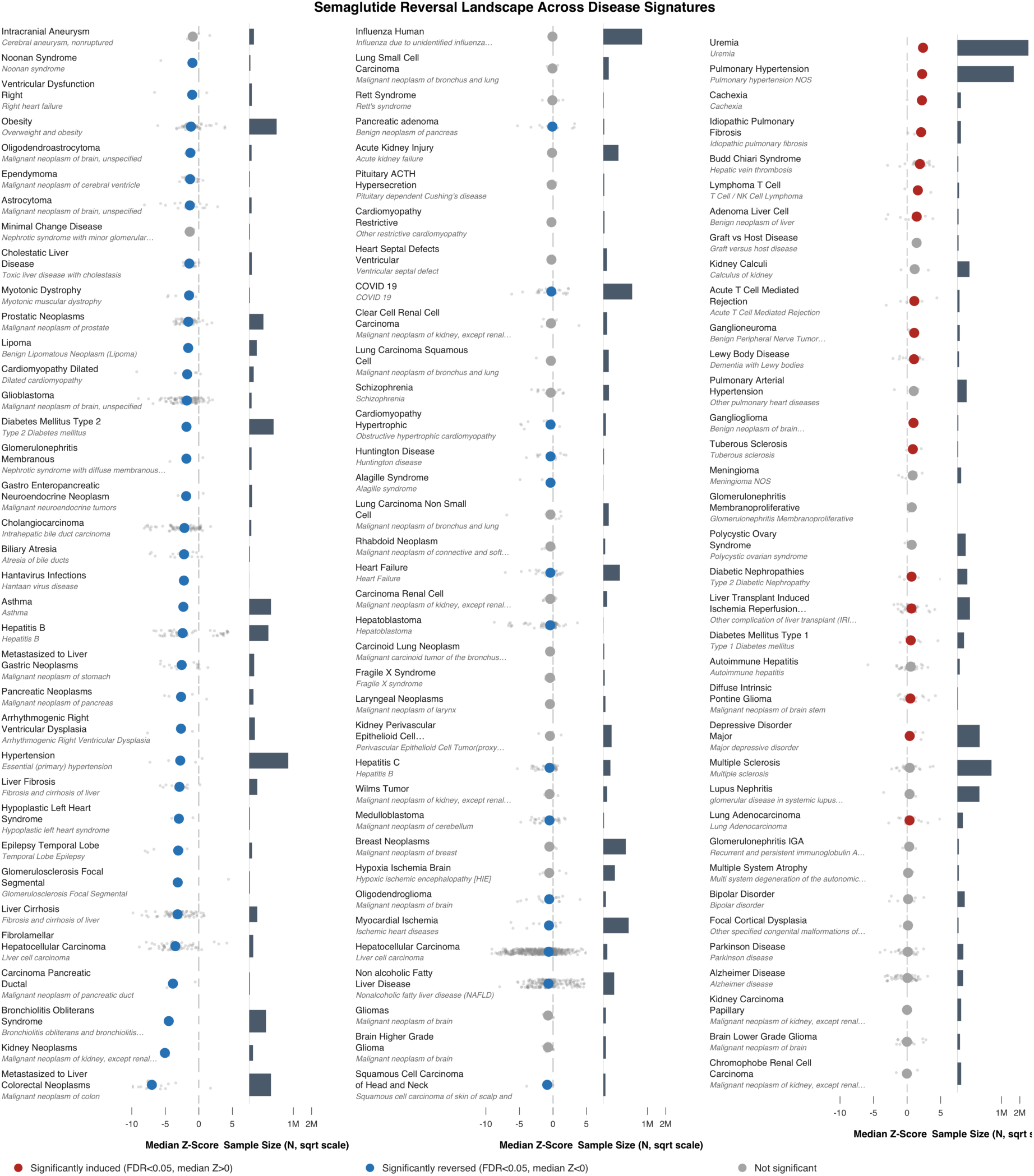
Semaglutide indication landscape characterized by transcriptomic reversal. For each disease, the dot plot displays the distribution of Z-RGES values across tissue-matched drug-disease comparisons, ordered by median Z-RGES and pooled across tissues. Individual dots represent tissue-level signature comparisons and large dots indicate the median Z-RGES per disease. Blue denotes diseases with statistically significant transcriptomic reversal, red denotes significant concordance (FDR < 0.05), and grey denotes non-significant comparisons. The adjacent bar plot shows total disease prevalence in the Mount Sinai Health System database on a square root scale, facilitating visualization across both common and rare diseases. For reversal without signatures pooled across tissue, see fig. S4B.

To better assess the full breadth of the predictive landscape, we performed literature-based validation with evidence-level stratification (table S8). For each disease, a literature search was undertaken to determine if it had a prior published association with semaglutide. If it existed, the nature of this association: therapeutic benefit, no effect, or contraindicated, was recorded, as well as the level of evidence, ranging from FDA approval to mouse studies. Predictive accuracy was assessed across evidence tiers: at higher levels of evidence such as FDA approval and meta-analysis, the framework performed best, predicting correctly for four out of five FDA approved indications, the exception being diabetic nephropathies (Fig. 4A).

**Fig. 4.**
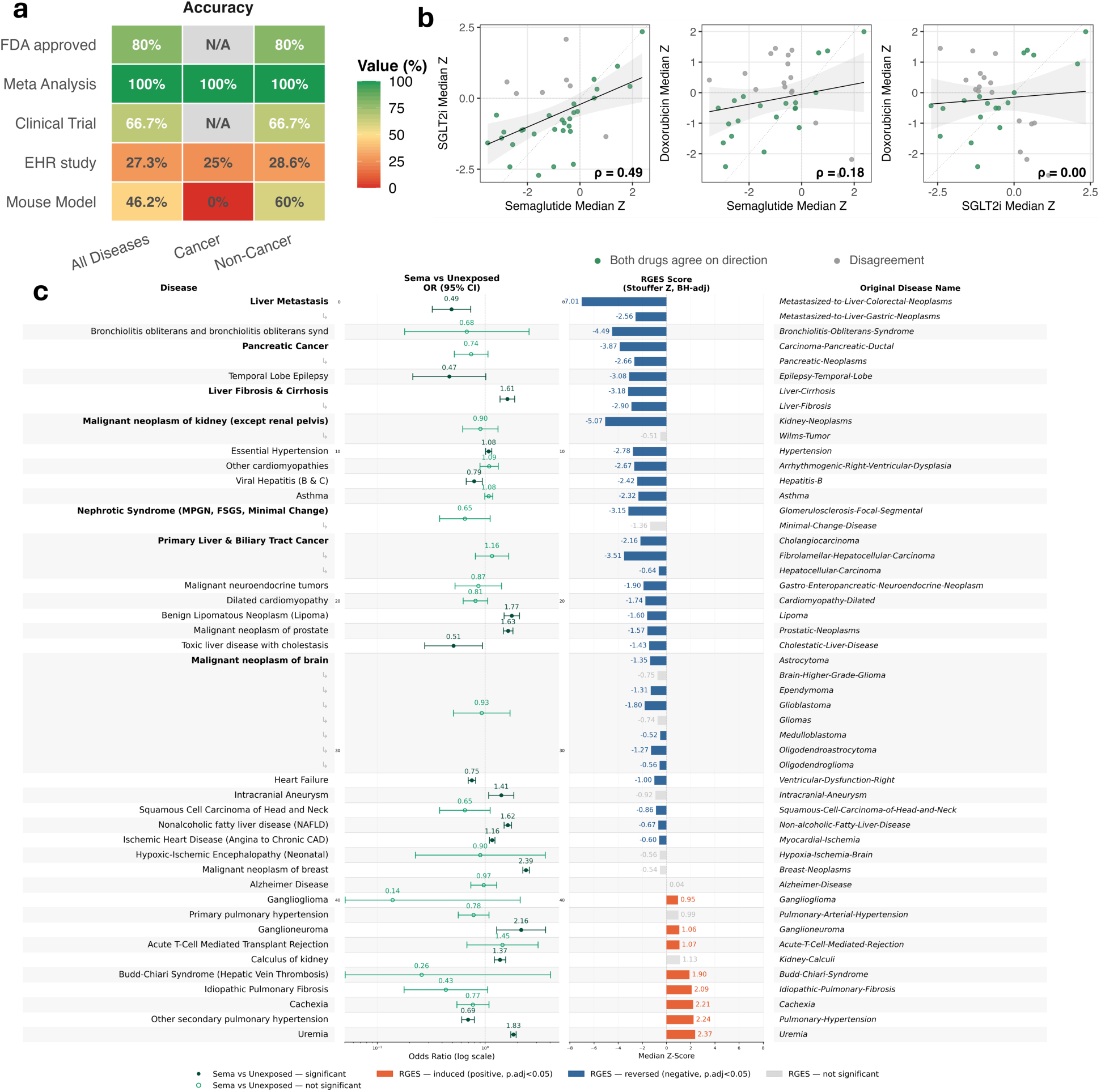
Validation of the framework. **(A)** Heatmap of semaglutide reversal method performance by evidence level (FDA-approved, meta-analysis, clinical trial, EHR study, mouse models) and disease group (all diseases, cancer, non-cancer), showing accuracy. Color indicates performance from red (weak) to green (strong); grey indicates insufficient data. **(B)** Pairwise correlation of median Z-RGES scores across diseases for semaglutide versus SGLT2 inhibitors (ρ = 0.49), semaglutide versus doxorubicin (ρ = 0.18), and SGLT2 inhibitors versus doxorubicin (ρ = 0.002). Neither doxorubicin comparison was significant at the p > 0.05 level. Green points indicate directional agreement between drugs; grey indicates disagreement. The moderate correlation between therapeutically convergent drugs and insignificant correlation with doxorubicin supports the discriminative validity of the framework. **(C)** Integrated comparison of EHR-derived odds ratios and Z-RGES scores across evaluated diseases. Left: odds ratios with 95% confidence intervals (log scale) for semaglutide-exposed versus unexposed patients. Right: BH-adjusted Stouffer Z-RGES scores, with blue indicating significant reversal, orange indicating significant concordance, and grey indicating non-significant scores. The rightmost column provides corresponding original disease names from the transcriptomic atlas.

To evaluate the reproducibility and specificity of the framework, we selected two comparator drugs as positive and negative controls. SGLT2 inhibitors, as an anti-diabetic with a similar, if mild, weight-reduction profile (*44*), were chosen as the positive control. Doxorubicin, as a well-profiled chemotherapeutic with extensive *in vivo* mouse studies and unrelated indication profile, was chosen as the negative control. Z-RGES was computed for both drugs across shared tissues, and their predictive scores were compared with those of semaglutide (table S9). Semaglutide and SGLT2 inhibitors exhibited a moderate positive correlation in predictive scores (Spearman’s ρ = 0.49), although the magnitude of reversal for shared indications was generally weaker for SGLT2 inhibitors, possibly owing to more moderate weight loss observed in patients taking SGLT2 inhibitors relative to semaglutide (*45*). In contrast, the correlations between doxorubicin and either antidiabetic treatment were not significant (p > 0.05), supporting the specificity of the framework’s predictions (Fig. 4B).

### *In vivo* transcriptomic reversal is concordant with EHR-based clinical evidence

To assess top-ranked predictions using real-world data, we analyzed EHRs from the Mount Sinai Health System via two arms, semaglutide vs unexposed patients, and semaglutide vs comparator drug SGLT2 inhibitors (fig. S5). To enable comparison between reversal scores and clinical outcomes, disease terms represented as MeSH terminology in GEOMeta were mapped to ICD-10-CM codes, the diagnostic classification system widely used for healthcare billing and administrative reporting. Because the two disease ontologies differ substantially in granularity and scope, the mapping required careful curation (table S10), with one notable limitation being the lack of ICD-10-CM codes for many specific tumors. For instance, although our transcriptomic atlas contains distinct signatures for individual brain tumor entities, several of these had to be mapped to broader ICD-10-CM categories such as “malignant neoplasm of the brain” or other generalized tumor diagnoses. Chart review may help resolve some of these ambiguities, but such approaches are difficult to scale. This loss of diagnostic specificity inherent to ICD-10-CM coding can reduce the resolution of downstream EHR-based validation analyses and poses an additional challenge for identifying clinical evidence in rare diseases and disease subtypes. Consequently, true therapeutic signals may be diluted or obscured when transcriptomically distinct diseases are aggregated into broader diagnostic categories within real-world healthcare data.

In the primary analysis of semaglutide versus unexposed controls, the semaglutide exposure arm was associated with lower odds ratio (OR) of six disease outcomes (Fig. 4C) (table S11). Notable associations included temporal lobe epilepsy (OR = 0.47, 95% CI: 0.21-1.01), liver metastasis (OR = 0.49, 95% CI: 0.32-0.76), toxic liver disease with cholestasis (OR = 0.51, 95% CI: 0.27-0.94), and other secondary pulmonary hypertension (OR = 0.69, 95% CI: 0.61-0.79). Temporal lobe epilepsy is of particular interest as previous studies have suggested a potential therapeutic role for semaglutide in this condition (*46*). Its transcriptomic reversal, together with a reduced odds ratio in the EHR analysis, provides convergent molecular and epidemiological evidence supporting this association. To assess overall concordance between the two approaches, we calculated the Pearson correlation across the 14 disease pairs that were significant in both the transcriptomic and EHR analyses (Pearson r = 0.50, p = 0.06). Although the correlation did not reach conventional statistical significance, its positive direction indicates broad agreement between the two orthogonal analyses. The limited number of disease pairs available for comparison likely reduced statistical power, suggesting that larger datasets may enable a more definitive assessment of concordance.

A secondary analysis comparing semaglutide initiators with SGLT2 inhibitor initiators was performed to evaluate semaglutide’s effectiveness relative to another antidiabetic drug class. Consistent with the more pronounced disease-reversal signals observed in the transcriptomic analysis, semaglutide exhibited lower odds of all significant outcomes compared with SGLT2 inhibitors. Notable examples included dilated cardiomyopathy (OR = 0.23, 95% CI: 0.15-0.36), essential hypertension (OR = 0.64, 95% CI: 0.55-0.72), heart failure (OR = 0.59, 95% CI: 0.51-0.68), and other cardiomyopathies (OR = 0.26, 95% CI: 0.18-0.35) (fig. S6, table S12).

### *In vivo* transcriptomic reversal prioritizes candidate indications for rare diseases

Having validated transcriptomic predictions in common diseases with available clinical evidence, we next investigated rare diseases, where clinical evidence remains limited. Of 54 diseases with fewer than 5,000 cases in the Mount Sinai Health System, 25 were significantly reversed. A subset of these candidates were chosen for further evaluation using independent datasets and network analyses, including bronchiolitis obliterans syndrome, arrhythmogenic right ventricular dysplasia (ARVD), myotonic dystrophy, Huntington’s disease, focal segmental glomerulonephritis, and membranous glomerulonephritis (Fig. 5, fig. S7). To facilitate interpretation, we provide representative Gene Set Enrichment Analysis (GSEA)-style visualizations of transcriptomic reversal for drug-disease signature pairs, including ARVD, myotonic dystrophy, and Huntington’s disease (Fig. 5A-C), together with the leading-edge genes which drive reversal (table S9). For instance, in arrhythmogenic right ventricular dysplasia, Fibroblast Growth Factor 13 (FGF13) was one of the four leading-edge genes driving the reversal of the down-regulated disease genes. FGF13 encodes an intracellular fibroblast growth factor homologous factor that directly binds and modulates the cardiac voltage-gated sodium channel and is essential for normal cardiac conduction, with loss of function increasing arrhythmia susceptibility (*47, 48*).

**Fig. 5.**
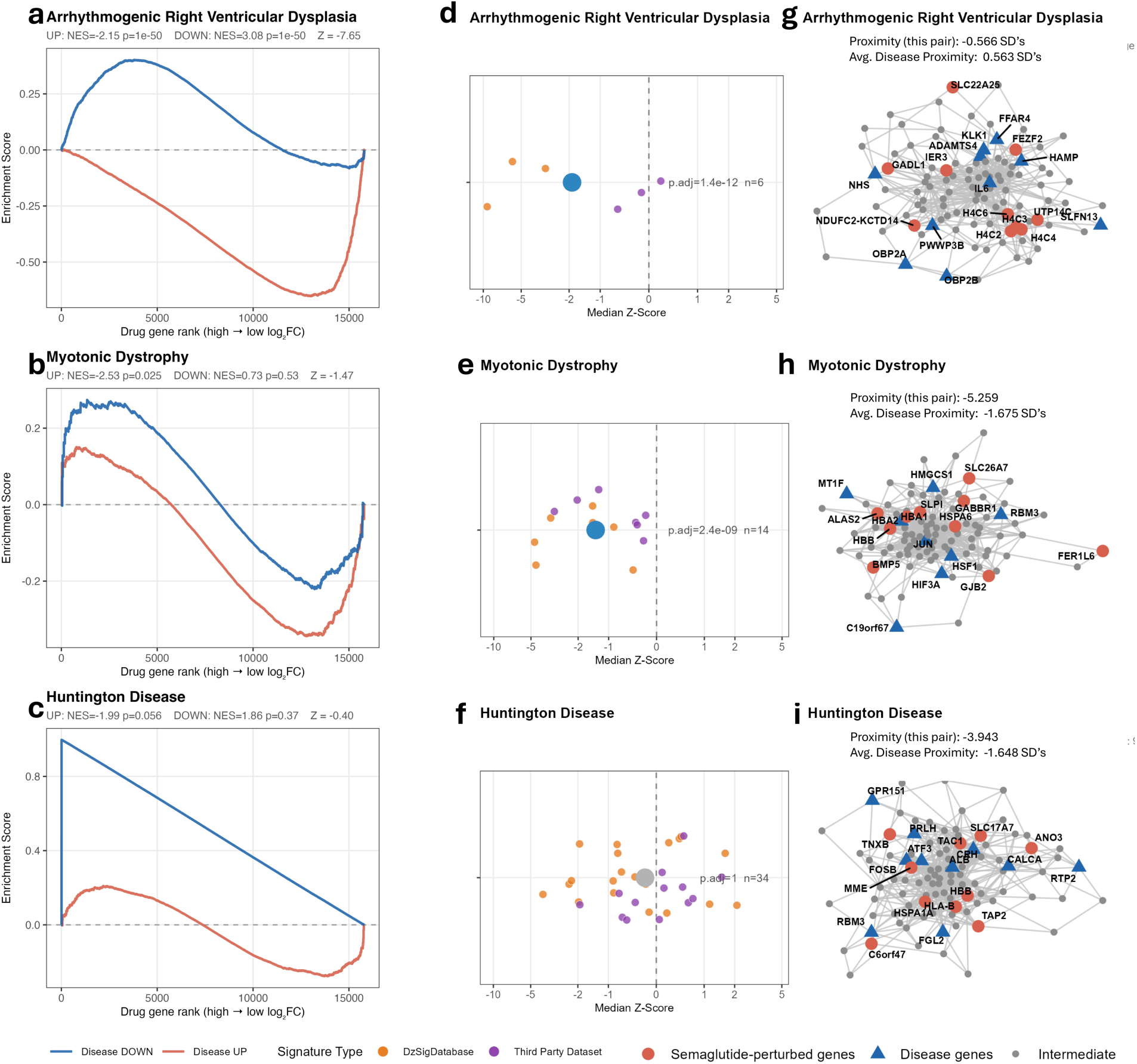
Rare disease validation and exploration. (A, B, **C)** GSEA-style enrichment plots showing transcriptomic reversal of disease gene expression signatures by semaglutide for ARVD (**A**) myotonic dystrophy (**B**) and Huntington’s disease (**C**). The blue curve traces enrichment of disease downregulated genes and the red curve traces enrichment of disease upregulated genes across the semaglutide-perturbed gene ranking (high to low log2 fold change). Crossing of the curves toward opposing ends of the ranked list indicates transcriptomic reversal. NES and Stouffer Z values are shown for each disease. **(D, E, F)** Distribution of Z-RGES scores across tissue-matched drug-disease comparisons for ARVD (**D**) myotonic dystrophy (**E**) and Huntington’s disease (**F**). Orange points represent comparisons from the harmonized disease atlas; purple points represent independent third-party disease signatures. The large central dot indicates the median Z-score. The Huntington’s disease panel (**F**) becomes non-significant upon inclusion of the full complement of available comparisons and is reported here for completeness given clinical interest in this indication. **(G, H, I)** Protein-protein interaction network proximity plots for ARVD (**G**) myotonic dystrophy (**H**) and Huntington’s disease (**I**). For each disease, the drug signature yielding the minimum proximity score was selected as the graphed pair, but median proximities are also displayed. Red circles represent semaglutide-perturbed genes, blue triangles represent disease-associated genes, and grey nodes represent intermediate network connections.

Because many rare diseases had only one qualifying disease signature in GEOMeta, we manually searched for additional signature sources for these diseases, and incorporated those which could provide reasonably robust signatures (Fig. 5D-F). Only bronchiolitis obliterans syndrome, arrhythmogenic right ventricular dysplasia, and myotonic dystrophy remained significantly reversed following the addition of the external datasets. Performing over-representation analysis using GO:BP pathways on the leading edge revealed significant pathways such as regulation of inflammatory response (raw p=0.002) for ARVD’s downregulated query set, or regulation of immune response (raw p= 0.009) in the case of myotonic dystrophy (table S13), a reasonable basis for biological reversal given ARVD’s promotion of inflammation (*49*) and myotonic dystrophy’s relation to immune dysfunction (*50*) . While repurposing a weight-loss drug to treat a genetic muscle-wasting disease may seem counterintuitive, the consistent and robust nature of the reversal of myotonic dystrophy type 1 in signatures from the heart and brain deserves further investigation. A hallmark of the disease is insulin resistance (*51*), as well as a dysregulation of mitochondrial processes and oxidative phosphorylation (*52*). That a similar drug, which might act without the promotion of weight-loss, could possibly mitigate symptoms has been proposed for similar diseases (*53*).

As a final mode of investigation, we performed network proximity analysis (Fig. 5G-I), an increasingly capable mode of computational drug discovery (*54, 55*). Because many of semaglutide’s most consequential effects are second-order and topologically distant from the GLP-1 receptor itself, we asked whether semaglutide’s transcriptionally perturbed genes sit more closely, within the human protein-protein interaction network, to each disease’s own perturbed genes than would be expected by chance. For each disease, semaglutide’s top ten DE genes (FDR < 0.05, ranked by absolute log2 fold change) were compared against the top ten DE genes of the disease’s reference signature(s). For each comparison we computed a symmetric network proximity score (the closest-path distance between gene sets, averaged in both directions) and tested its significance against a degree-preserved null generated by 1,000 randomized gene sets matched to the size and connectivity of the true sets, yielding a Z-score in which more negative values indicate proximity closer than chance.

Of the six diseases examined, myotonic dystrophy showed the most consistent evidence of non-random proximity: 5 of 7 independent comparisons trended proximal, with a mean z of −1.68 (FDR = 1.61 × 10⁻^5^). The most mechanistically suggestive direct interaction was between HSF1 and HSPA6, linking semaglutide-perturbed genes to the heat-shock proteostasis machinery; although myotonic dystrophy pathology is driven principally by RNA toxicity(*56*), heat-shock proteins have independently been implicated in the disease, most directly the small heat-shock protein HspB2/MKBP, which binds and stabilizes DMPK (*57*), making proteostasis a plausible, if indirect, mechanistic axis for this connection.

Huntington’s disease showed an unexpectedly strong and consistent proximity signal: 20 of 21 independent comparisons were proximal, with a mean z of −1.65 (FDR = 1.61 × 10⁻^5^) and six comparisons individually exceeded z ≤ −2. This dissociation between an absence of transcriptional reversal and strong network-level proximity suggests semaglutide’s targets may sit mechanistically adjacent to the Huntington’s disease module without producing the compensatory expression changes captured by reversal scoring. We therefore do not regard semaglutide as a lead candidate in Huntington’s; rather, this is an open, mechanistically ambiguous signal warranting targeted follow-up.

By contrast, ARVD did not hold up under randomization: mean z was positive (+0.56, FDR = 8.35 × 10⁻^1^), and only one of three available comparisons trended toward proximity. Despite the lack of proximity, we note that this comparison was only between the top ten DE genes, while reversal included query sets of up to 500. The consistent reversal may still be a signal of broader systemic effects that are not captured by local network proximity.

## DISCUSSION

Despite substantial advances in drug discovery, de novo drug development remains highly inefficient, typically requiring 10**-**17 years and more than 2 billion USD to bring a new chemical entity to market (*58, 59*). Indication expansion has become a necessary strategy in drug discovery owing to this gap; notable examples including Sildenafil, initially developed for angina and later approved for erectile dysfunction (*60*); Thalidomide, originally developed for morning sickness and subsequently repurposed for multiple myeloma(*61*); and Ibrutinib, first investigated in mantle cell lymphoma and later established as a standard-of-care treatment for chronic lymphocytic leukemia (*62*). While numerous approaches have been explored to identify new indications, most have relied heavily on domain expertise, serendipitous clinical observations, or mechanistic insights. For example, the recent clinical success of semaglutide in obesity emerged largely from observations made during clinical studies rather than systematic indication screening (*63*). Such strategies are often most effective in common diseases, where large patient populations facilitate the detection and validation of unexpected therapeutic effects (*64*). In contrast, identifying new indications for rare and uncommon diseases remains challenging because of limited patient numbers and sparse clinical evidence. This work addresses this gap by leveraging the transcriptomic reversal paradigm, tissue-specific drug signatures, and a large-scale compendium of human disease signatures to systematically evaluate indication expansion opportunities across diverse diseases and tissues. This framework, available on our OCTAD 2.0 portal: https://apps.octad.org/octad2, enables unbiased, data-driven prioritization of potential therapeutic indications, including diseases for which conventional clinical observation-based approaches may be impractical.

Fundamentally, transcriptomic reversal seeks to determine whether a drug-induced phenotype opposes a disease-associated phenotype. By representing disease and drug in this way, screening at scale becomes possible with available transcriptomic signatures and an appropriate reversal scoring function. Until now, however, drug-induced signatures have been derived predominantly from immortalized cell lines, whose scalability comes at the expense of biological fidelity. Many drugs exert pleiotropic effects across multiple tissues through interconnected physiological and molecular processes, including well-established inter-organ communication pathways such as the gut–brain, liver–brain, and gut–liver axes. Consequently, reversal analyses that do not account for systemic effects may provide an incomplete or inaccurate representation of a drug’s therapeutic potential. By combining harmonized transcriptomic expression matrices and standardized sample metadata created by an agentic large-language model-based (OpenAI’s GPT-5) annotation pipeline, we can obtain *in vivo* signatures of drugs and diseases at scale. This resource enables a more system-aware implementation of transcriptomic reversal by incorporating organism-level drug responses while the standardized signature generation pipeline and Z-transformed RGES facilitate direct comparison of reversal scores across diseases. The resulting framework successfully recovered known semaglutide indications, identified emerging therapeutic opportunities, and demonstrated concordance with real-world clinical outcomes.

Moreover, this framework may serve as a scalable exploratory tool throughout the drug discovery pipeline, not merely for the repurposing of established drugs as we sought to demonstrate here. Because multi-tissue RNA-seq profiling can be performed at a fraction of the cost of late-stage clinical development, systemic reversal screening could be incorporated early in drug development to identify potential liabilities and therapeutic opportunities before substantial resources are committed. Notably, because the framework relies solely on drug-induced multi-tissue transcriptomic profiles, it is not limited to small-molecule therapeutics. In principle, it can be applied to a wide range of therapeutic modalities, including biologics, antibodies, probiotics, dietary supplements, and cell-based therapies, as long as their *in vivo* molecular responses can be profiled. Given that approximately 90% of drug candidates entering clinical trials ultimately fail to achieve regulatory approval (*22*), early assessment of tissue-wide reversal patterns may help prioritize indications with stronger biological support, de-risk clinical development, and reveal alternative disease areas in which a compound may have greater therapeutic potential. In the case of semaglutide, the lack of reversal for neural diseases like Parkinson’s, major depressive disorder, and Alzheimer’s could have been useful prior to the initiation of expensive GLP-1 clinical trials for these diseases (*41–43*).

While transcriptomic reversal is not the only large-scale screening framework for drug discovery, it differs from current drug repurposing approaches that operate within a mechanism-first paradigm: one gene, one target, one disease (*54*). Network-based in silico drug screening is one approach which formalizes this logic, scoring drug-disease pairs by their network distance from the drug target, an approach with solid grounding, but inherently less sensitive to off-target and downstream effects. For drugs like semaglutide, whose most clinically consequential effects are second-order effects (*24, 65*), this framework would miss new indications using the traditional approaches. Conversely, transcriptomic reversal captures these system-level responses but does not, by itself, establish whether the reversed genes are mechanistically related to disease biology. Two signatures may exhibit some transcriptional opposition despite involving only tangentially related pathways. For those drugs whose effects are cell-autonomous and receptor-proximal, the most meaningful computational screen would include both classical proximity and reversal screening, capturing opposing phenotypes and proximal drug/disease pairs with their orthogonal approaches providing separate perspectives (*55*). For drugs with broader systemic effects, we adapted the proximity framework by assessing whether genes most strongly perturbed by the drug are themselves proximal to disease-associated networks, thereby providing an orthogonal filter for biological plausibility. Although further exploration of this integrated framework is warranted, the ability to combine transcriptomic reversal with network-based mechanistic evidence directly from RNA-seq and existing systems-biology data may offer a scalable strategy for prioritizing therapeutic candidates and reducing translational risk early in drug development. However, the addition of proximity screening is not required for our framework.

Some limitations of this study should be acknowledged. First, disease pathogenesis and drug mechanisms of action are highly complex, and transcriptomic profiles alone may not fully capture the underlying biology. Consequently, some true drug-disease relationships may not be identified by a transcriptome-based reversal framework, as was the case here with diabetic nephropathies. In addition, the current implementation relies on bulk RNA-seq data, which lacks the cellular resolution afforded by single-cell and spatial transcriptomic technologies. As higher-resolution and multi-modal profiling datasets become increasingly available, the framework could be extended to incorporate these data modalities and provide a more comprehensive representation of disease and drug states. Second, the semaglutide-induced transcriptomic signatures used in this study were derived from rodent models, and their translatability to human biology cannot be assumed. These rodents were on a high fat diet, and since the theoretical best use of transcriptional reversal would involve the comparison of a given disease and a drug signature derived from a model undergoing that same disease, to capture disease-specific effects of a drug, the most technically robust reversal here is that of obesity. Such experiments are not practical to scale; nevertheless, our framework could provide an initial set of diseases to pursue further. The transcriptomic and EHR analyses provide complementary but distinct perspectives on therapeutic effects: transcriptomic reversal evaluates the extent to which a drug counteracts an established disease-associated molecular state, whereas the EHR analysis primarily assesses associations with disease incidence in a preventative setting. Although these approaches help address each other’s limitations, they should not be interpreted as equivalent measures of treatment efficacy. Additional limitations apply to the EHR analysis. The reported odds ratios were derived from matched cohorts and represent unadjusted estimates. In the absence of covariate-adjusted survival analyses or causal inference frameworks, these findings should be interpreted as associative rather than causal. Statistical power was also limited for rare outcomes with fewer than ten incident cases, increasing uncertainty in effect estimates. Finally, and most importantly, all findings presented here should be considered hypothesis-generating. Experimental validation in relevant biological systems and prospective clinical studies will be required before any predicted indication-expansion opportunity can be considered established.

## MATERIALS AND METHODS

### Study Design

We sought to develop and validate a framework for systematic indication expansion that compares *in vivo* drug and disease signatures, for an accurate reflection of the true transcriptional states that drug and disease perturbations cause in the mammalian organism. The hypothesis was such that *in vivo* multi-tissue drug signatures would reverse human disease signatures in a tissue-context-dependent manner, and enrich for clinically relevant indications. Semaglutide was chosen as the primary case study, with the rare disease analyses being merely exploratory and created after inspection of the primary reversal results.

Three components composed this study: a controlled laboratory experiment using rats, a computational analysis of human disease signatures, and a retrospective observational analysis of electronic health records.

16 male Dahl salt-sensitive rats 4 weeks of age were randomly assigned to control (10% kcal fat) or high-fat (60% kcal fat) diet. This was maintained for 21 weeks before they were randomized a second time to receive semaglutide (0.1 mg/kg/day, subcutaneous) or vehicle for 28 consecutive days. Injections were given at a fixed daily time to limit circadian variability.

Tissues were collected from seven organs 28 days after treatment initiation. An RNA integrity number below 5 was defined as an exclusion criterion before sequencing; on this basis, most pancreas and colon samples failed quality control and both tissues were excluded from all subsequent analyses. Sampling replicates consisted of individual animals within each diet– treatment group, and experimental replicates consisted of the retained tissues profiled per animal. To assess generalizability across species, dosing regimens, and durations, the rat signatures were supplemented with 16 publicly available semaglutide-perturbed mouse datasets, yielding 35 drug signatures in total.

The primary endpoint was the reversal of human disease signatures by an *in vivo* drug signature, quantified as a permutation-based Z-score of the reversal gene expression score. The secondary endpoint was the association between semaglutide exposure and incident disease in electronic health records. Both were specified before the EHR analysis was conducted. Because many diseases were tested, all *P* values were corrected for multiple comparisons using the Benjamini– Hochberg procedure within each comparison, with significance defined as FDR < 0.05.

The EHR component was a retrospective observational analysis and was not randomized. Two parallel comparisons were pre-specified: semaglutide versus drug-unexposed patients, and an active-comparator analysis of semaglutide versus SGLT2 inhibitors. A new-user, intention-to-treat design was applied with eligibility criteria defined before analysis, and confounding was addressed by propensity score matching rather than randomization. The study period and data cutoff were fixed in advance.

### Construction of the Human Disease Signature Atlas

To generate a reference landscape of human disease-associated transcriptional phenotypes, publicly available human bulk RNA-sequencing datasets were aggregated from the Gene Expression Omnibus (GEO) (*25*), Sequence Read Archive (SRA) (*66*), The Cancer Genome Atlas (TCGA) (*67*), and Treehouse pediatric repositories (*33, 34*). Gene-level expression matrices were primarily obtained from the ARCHS4 (*26*) resource, which provides uniformly processed RNA-seq quantifications generated using a standardized alignment and quantification workflow. Datasets not represented within ARCHS4 were processed using compatible transcript quantification pipelines to ensure consistency across studies (*68*). All expression profiles were harmonized to a common GRCh38 gene annotation space prior to downstream analyses. Sample-level quality control was performed before disease signature generation. Quality metrics included total mapped reads (>10 million) and number of detected genes (>10,000). Principal component analysis (PCA) was performed separately within each study using the top 5,000 most variable genes, scaled to unit variance following CPM normalization and log2 transformation, to identify transcriptomic outliers. To account for sample size constraints, the first min (10, number of samples − 1) principal components (PCs) were retained per study. Studies with fewer than 6 samples were exempted from PCA-based outlier filtering, as covariance structure cannot be reliably estimated below this size, but remained subject to the read-depth and gene-detection QC criteria described above. For studies with at least 6 samples, a sample was classified as an outlier if its 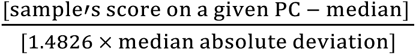 exceeded 5 on any retained PC, and was excluded from downstream analyses. Duplicate samples identified across repositories were removed to prevent redundancy. Following quality control, a large-scale transcriptomic compendium spanning ∼600,000 samples, 60,000 genes, 50+ tissues, and 3000+ studies was assembled for downstream analyses.

### Metadata Harmonization and Disease Annotation

Public transcriptomic repositories frequently contain incomplete, inconsistent, and heterogeneous sample metadata, limiting large-scale disease integration. To address this challenge, we used GEOMeta (*31*), an agentic LLM (OpenAI’s GPT-5)-guided metadata extraction and harmonization framework developed in our group for large-scale GEO sample annotation. The development of GEOMeta is detailed elsewhere (*31*). Briefly, GEOMeta retrieves GSE- and GSM-level metadata from user-provided GEO Series identifiers and uses study-level and sample-level information, including series titles, summaries, overall design fields, sample titles, source names, characteristics fields, and free-text descriptions, to generate structured sample annotations. GEOMeta generated standardized annotations including disease status, RNA source, tissue source, control or reference status, treatment or perturbation status, and experimental context. Raw annotations were standardized using field-specific post-processing rules, conservative derived-field inference, and ontology- or resource-based mapping. Disease labels were normalized to controlled disease terms using reviewed disease mappings together with CTD/MEDIC disease names and synonyms to reduce redundancy from inconsistent disease nomenclature. Ambiguous or unresolved disease assignments were routed to manual review rather than automatically assigned. GEOMeta annotation quality was assessed using layered quality-control and validation procedures, including manual benchmark evaluation, text-evidence validation against original GEO metadata, and external concordance analyses with DiSignAtlas for disease and tissue annotations. Using this framework, nearly 600,000 publicly available transcriptomic samples were harmonized into a standardized sample-level annotation schema. Only studies with adequate metadata quality, clearly defined disease phenotypes, and biologically interpretable experimental designs were retained for disease signature generation. One additional round of manual review of all disease-tagged samples was performed to ascertain accuracy of annotations.

### Selection of Surrogate Controls

A substantial proportion (62%, 1885/3026 *in vivo* studies) of publicly available disease transcriptomic studies lack appropriate healthy control samples, particularly for rare diseases and surgically inaccessible tissues. To address this limitation, surrogate controls were identified using a previously published latent transcriptomic embedding framework(*69*). A fully connected autoencoder was trained using healthy GTEx samples together with high-confidence ARCHS4 samples possessing reliable tissue annotations. Prior to training, expression matrices were normalized and log-transformed. The network compressed high-dimensional transcriptomic profiles into a 64-dimensional latent representation optimized using mean squared reconstruction loss (64 encoded features, 128 batch size, 100 epochs, 0.0002 learning rate). Following training, disease samples and GTEx controls were projected into the latent space. Similarity between samples was quantified using Pearson correlation, Spearman correlation, and cosine similarity metrics calculated on latent embeddings. For studies lacking native controls, the highest-ranking tissue-matched GTEx samples were selected as surrogate controls. Studies for which biologically plausible control matches could not be identified (correlation < 0.5) were excluded from downstream analyses.

### Differential Gene Expression Analysis

Disease-specific transcriptomic signatures were generated independently for each study using edgeR (v4) within a negative binomial generalized linear modeling framework. Library composition bias was corrected using trimmed mean of M-values (TMM) normalization. Genes were retained if they had counts-per-million (CPM) > 1 in at least 2 samples. For studies containing native matched controls, disease status was modeled as the primary contrast variable. For studies utilizing surrogate controls, unwanted technical variation was estimated using RUVseq and incorporated into the design matrix as covariates (*70*). Dispersion estimates were calculated using empirical Bayes methods implemented within edgeR, including common, trended, and tagwise dispersions. Differential expression significance was assessed using quasi-likelihood F-tests(*71*). Multiple testing correction was performed using the Benjamini-Hochberg procedure, and genes with an adjusted FDR below 0.05 were considered statistically significant. For each study, standardized disease signatures consisting of log2 fold changes, average logCPM, raw p-values, adjusted p-values, and test statistics were generated.

### Validation of Disease Signatures

To evaluate the biological validity and reproducibility of the generated disease signatures, we compared our results against independent transcriptomic disease resources and curated disease-gene databases. First, signatures were compared against corresponding signatures from DiSignAtlas, a large-scale disease transcriptomics resource derived from bulk and single-cell gene expression studies. For diseases represented in both resources, pairwise concordance was assessed using Spearman correlation between gene-level log2 fold changes. Correlations were calculated using the intersection of genes present in both signatures. High correlation coefficients were interpreted as evidence that the generated signatures recapitulated independently derived disease-associated transcriptional phenotypes.

Second, pathway-level validation was performed via GSEA using Hallmark, Reactome, and GO:BP gene sets obtained from MSigDB. Concordance between enriched pathways and generally known disease mechanisms was manually evaluated for representative diseases (table S1). Together, agreement with DiSignAtlas and recovery of expected biological pathways provided orthogonal validation of disease signature quality prior to downstream transcriptomic reversal analyses.

### Disease Signature Quality Control and Tissue Matching

To ensure biologically meaningful reversal analyses, disease signatures underwent additional quality-control procedures following aggregation. Signature stability was assessed through cross-study concordance analyses, and diseases exhibiting inconsistent transcriptional directionality across studies were flagged for manual review.

Disease signatures were grouped according to tissue source to enable biologically matched transcriptomic reversal analyses. Only tissue-matched comparisons were permitted during reversal analysis. For example, semaglutide-perturbed liver signatures were compared exclusively against liver-derived disease signatures, semaglutide-treated heart signatures against heart-derived disease signatures, and similarly for kidney, lung, and brain tissues. For those diseases with signatures from multiple tissues, we treated them separately.

Disease query gene sets used for reversal analysis consisted of significantly dysregulated genes meeting thresholds of absolute log2 fold change > 1 and FDR < 0.05. Following quality control, validation, aggregation, and tissue matching, disease signatures representing 108 unique diseases were retained for downstream transcriptomic reversal analysis. The final atlas consisted of over 300 unique diseases consisting of 979 individual signatures, spanning multiple tissues and organ systems, from which 108 diseases possessed tissue-matched counterparts suitable for transcriptomic reversal analysis against semaglutide-perturbed signatures.

### Web Portal Development and Deployment

To enable interactive exploration of the disease signature atlas, a web portal OCTAD 2.0 https://apps.octad.org/octad2/ was developed using the R Shiny framework. The Disease Explorer module supports visualization of disease-level differential expression (volcano plots, ranked DE genes), gene queries across all signatures, disease-disease correlation, and pathway enrichment (GO, KEGG, Reactome, WikiPathways, Hallmark); diseases with multiple datasets are consolidated via random-effects meta-analysis. The Drug Explorer module supports two query directions: a Drug Repurposing tool that scores LINCS compounds for their ability to reverse a disease signature (RGES-based), and an Indication Expansion tool that takes a user-supplied gene signature (up/down gene list, optionally tissue-restricted) and screens all disease signatures for significant reversal or concordance via hypergeometric enrichment testing, enabling indication-expansion analysis for any drug or novel compound signature.

### Drug Signature Generation and Characterization

#### Rat treatment design

Sixteen male Dahl salt-sensitive (SS) rats, bred in-house at Michigan State University, were used in this study. At 4 weeks of age, animals were randomized to one of two dietary conditions: a control diet (CD; 10% kcal from fat) or a high-fat diet (HFD; 60% kcal from fat). Diet was maintained for 21 weeks prior to the initiation of drug treatment, and was sustained throughout the treatment period. Following the dietary acclimation period, animals were randomized to receive either semaglutide or vehicle. Semaglutide free base (Item #40231; Cayman Chemical Company, Inc., Ann Arbor, MI) was administered subcutaneously at a dose of 0.1 mg/kg/day for 28 consecutive days. Vehicle-treated animals received an equivalent volume of a solution composed of approximately 1% DMSO, 1% Tween 80, and 98% USP saline. All injections were administered at the same time each day to minimize circadian variability.

All procedures described in this study have been reviewed and approved by the Michigan State University Institutional Animal Care and Use Committee (IACUC) and have been conducted in accordance with the Guide for the Care and Use of Laboratory Animals and applicable AVMA guidelines. The proposed studies are covered under MSU IACUC Protocol #PROTO202400255

#### RNA Extraction and Quality Control

Tissues from the high-fat diet cohort were harvested 28 days post treatment start. We chose major organs including pancreas, liver, kidney, lung, heart, colon, and brain for RNA library preparation. Tissues were homogenized in TRIzol reagent using a Fisherbrand™ 850 Homogenizer (Thermo Fisher Scientific) at 4°C. Each sample was homogenized for three 15-second intervals with brief pauses between cycles to ensure complete tissue disruption. Total RNA was extracted using the RNeasy Mini Kit (Qiagen) according to the manufacturer’s instructions. To remove residual genomic DNA, an on-column DNase I digestion was performed during RNA purification using the RNase-Free DNase Set (Qiagen) according to the manufacturer’s instructions. RNA concentration was measured using a NanoDrop spectrophotometer. RNA quality and integrity were assessed by agarose gel electrophoresis and an Agilent 2100 Bioanalyzer. Samples with an RNA integrity number (RIN) < 5 were excluded from further analysis. As a result, most samples from pancreas and colon did not pass quality control, therefore, we excluded pancreas and colon in the following experiments. Qualified RNA samples were submitted to Novogene Co., Ltd. for RNA-seq library preparation and high-throughput sequencing.

RNA-seq was performed by Novogene. Raw reads were aligned to the GRCr8 reference genome using STAR and quantified to generate gene-level raw counts using RSEM.

#### Data Processing

Semaglutide-perturbed mouse datasets (N = 16) from GEO were used to augment the analysis. Raw counts were retrieved from ARCHS4, and metadata was manually procured (table S2). Signatures were generated based on tissue source, diet, dosage, and duration (N = 35) using DESeq2 version 1.50.2. Genes were mapped across species orthologs using Ensembl BioMart. Genes provided as Ensembl IDs were matched directly on Ensembl accession. Where a gene mapped to multiple orthologs, each ortholog was retained as an independent entry carrying that gene’s expression statistics. Where multiple genes mapped to the same ortholog, their log2 fold changes were collapsed by taking the median at the point of ranked-signature construction for connectivity scoring. Genes without an identifiable ortholog were excluded from scoring.

#### Gene Set Enrichment Analysis of Drug signatures

GSEA was performed using the fgsea R package on Hallmark, Reactome, and GO:BP gene sets sourced from MSigDB via the msigdbr R package. Genes were ranked by log2 fold change, and gene sets were considered significantly enriched at an adjusted p-value < 0.05. Signatures were grouped into cohorts: all signatures, mice, rats, and each tissue with enough contributing signatures (n >3) being liver, brain, white adipose tissue, and heart. Within each cohort, we combined per-signature evidence for each pathway using a signed Stouffer method 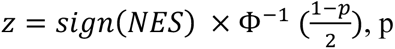, p floored at 10⁻¹⁰; 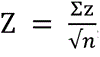, Benjamini-Hochberg corrected within every particular combination of cohort and pathway database. A pathway was conserved in a cohort if it had ≥3 contributing signatures, directional consistency ≥0.90 (pooled core) or ≥0.80 (individual cohorts), FDR < 0.05, and |median NES| ≥1.0.

Cohort coherence was confirmed by comparing within-versus between-tissue pairwise Spearman correlations of NES profiles. This was tested with one-sided Wilcoxon tests, required median margin ≥0.10 and p < 0.05. Tissue specificity was assessed in two ways: a strict filter, which is to say, conserved in the tissue, but non-significant and |NES| < 1.0 in ≥ 90% of other signatures, and a relative-enrichment test: conserved in the tissue and significantly higher there than elsewhere (FDR < 0.05, median-NES margin ≥0.75). Directional discordance, meanwhile, was assessed between tissues. Discordance required a pathway to be independently significant in two tissue sets (FDR < 0.05), in the opposite direction, and |median NES| ≥1.0 in each; this was designated claim-grade only when both cohorts had ≥3 signatures and were coherent.

Every claim was validated by leave-one-out: each contributing signature was removed in turn (both cohorts for discordance), the claim’s criteria recomputed on the reduced set, and the claim called robust only if it held across all iterations. Analyses used R (tidyverse, fgsea, DESeq2).

#### Cell-type Deconvolution of Bulk Data

Deconvolution of the cell types present in the signature was performed using sigatlas (*72*), an in-house bulk RNA-seq curation pipeline. For the mice signatures, the default cell-type markers offered by sigatlas were used. For the rat signatures, custom markers for brain (*73*), heart (*74*), liver and kidney (*75*), and lung (*76*) were assembled from literature (table S14). Exact sigatlas output can be found in Supplementary Tables 15 and 16.

#### Reversal of Disease-associated signatures

We quantified the reversal of disease gene expression via Z-RGES. Drug and disease signatures were paired for reversal on the basis of the source tissue, e.g. semaglutide-perturbed lung signatures were reversed against disease signatures sourced from human lungs. 108 unique diseases with matched tissue sources were available from the wider 300 disease database. Drug signatures were used as the ranked set, and the disease signatures the source of the query sets. Query gene sets consisted only of those genes with log2 fold change > 1 and FDR < 0.05, and were capped at 500 genes. Raw connectivity scores were computed using a Kolmogorov-Smirnov-based scoring function. To normalize these scores and enable comparison despite differences in gene universe across signatures, for a given drug disease pair, an empirical null distribution was generated for each pair by randomly sampling gene sets of equivalent size from the gene universe (300 permutations). This was repeated across 5 independent random seeds, yielding 5 Z-scores per pair; the final reported score is the median Z-score across these 5 iterations, providing a robust estimate of connectivity relative to the random distribution. Diseases were then ranked by the median Z-score across all comparisons for a given disease, and the significance of this median disease Z-score computed via BH-adjusted Stouffer’s method (FDR < 0.05).

#### Network Analysis

The package string_db was used (version 12.0, species 9606 ‘human’, with a score threshold of 400 and a ‘functional’ network type) to construct the human interactome. For a given disease, semaglutide transcriptomic signature’s top ten DE genes was compared against the disease’s top-ten DE genes using a symmetric proximity measure, yielding multiple independent drug-disease comparisons per disease. Statistical significance for each comparison was assessed against a degree-preserved null distribution, following Guney et al. (*55*). Network nodes (n = 19,488) were partitioned into 20 quantile-based connectivity bins (∼974 nodes/bin). For each comparison, 1,000 random gene sets were generated by replacing each real gene with a randomly selected node from its same connectivity bin (sampled without replacement within a single draw), independently for the drug and disease gene sets; proximity was recomputed for each random draw using the identical measure applied to the observed data, yielding a null mean (μ_R) and standard deviation (σ_R). Observed proximity was expressed as 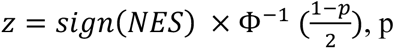, with more negative values indicating proximity closer than expected by chance. For diseases with more than one independent disease-reference signature, each drug’s Z-score was first averaged across its repeated disease-reference comparisons, so that non-independent repeats of the same drug were not treated as separate observations. Per-disease significance was then obtained by combining the resulting per-drug Z-scores via Stouffer’s method, 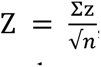, with resulting one-sided p-values (testing for proximity closer than chance) BH-adjusted across the diseases examined.

#### Secondary dataset validation

Signatures from GEO not in the disease atlas were sourced from GSE197307, GSE198321, GSE216841, GSE224858, GSE232649, GSE284565, and GSE290834. As GSE290834 is a single-cell dataset, pseudobulk signatures were constructed by summing gene counts across cell types across four cohorts, Graft-vs-Host-Disease bronchiolitis obliterans and bronchiolitis obliterans, between immune and non-immune cell type fractions.

### EHR-Based Pharmaco-Epidemiological Analysis

#### Data source

Data was obtained from the Mount Sinai Health System EHR, which contains structured clinical data (*77, 78*). A combination of ICD-10-CM codes and condition source value text extracted from an OMOP-structured condition occurrence table was used to determine diagnosis. The disease terms in GEOMeta were mapped to ICD-10-CM codes and clinical synonyms; diseases with no direct ICD-10-CM equivalent were mapped to their closest parent codes. table S10 provides a complete mapping of 82 disease concepts, including associated ICD-10-CM codes and search synonyms. EHR-based association analysis was performed on 50 diseases with sufficient transcriptomic evidence (top 40 most reversed, bottom 10 most induced). All Mount Sinai related analyses were conducted under an IRB-approved study.

#### Study population

Two parallel analyses were conducted: semaglutide versus drug-unexposed patients and an active comparator analysis of semaglutide versus SGLT2 inhibitors. The semaglutide-unexposed analysis comprised 55,302 semaglutide users and 320,805 patients with qualifying diagnoses, at least one healthcare encounter within the study period, and no prior or current Glucagon-like peptide-1 receptor agonists (GLP-1RA) exposure at any time. The semaglutide-SGLT2 inhibitor analysis included 67,212 patients with qualifying type 2 diabetes mellitus (E11.x) or obesity (E66.x) diagnoses identified from 96,659 patients with semaglutide or SGLT2 inhibitor exposure. The study period was January 1, 2018, through December 31, 2025, with a data cutoff of March 31, 2026.

#### Exposure definition and index date

Semaglutide exposure included all formulations (Ozempic, Wegovy, Rybelsus). SGLT2 inhibitor exposure included empagliflozin (Jardiance), dapagliflozin (Farxiga), canagliflozin (Invokana), and ertugliflozin (Steglatro), treated as a drug class. The index date was the date of first dispensing for drug-exposed cohorts and the earliest of the qualifying diagnosis date or first eligible encounter on or after study period for the unexposed cohort.

#### Eligibility criteria

An intention-to-treat approach (*79*) with new user design was applied, excluding patients with prior use of the assigned drug class within 365 days. At least two dispensing records were required to confirm continuous exposure. Additional eligibility criteria included age 18-85, at least one encounter in the preceding 12 months, and no prior type one diabetes mellitus (T1DM) or ESRD (end stage renal disease). Simultaneous initiation of both drug classes within seven days and prevalent outcomes at index were excluded. The patient selection workflow is presented in fig. S5, and the baseline characteristics are presented in table S17.

#### Propensity score (PS) matching

PS were estimated by logistic regression including age, sex, race, ethnicity, BMI, T2DM duration, T2DM and obesity flags, comorbidities (chronic kidney disease, cardiovascular disease, heart failure, atrial fibrillation, hypertension, dyslipidemia), prior metformin and DPP4 inhibitor use, and calendar year strata (2018–2019, 2020–2021, 2022–2023, 2024–2025). Missing BMI values were imputed using multiple imputation by chained equations (MICE) (*80*), with the nearest available BMI measurement to the index date used for each patient. The semaglutide-unexposed analysis used 1:2 matching, and the semaglutide-SGLT2 inhibitor analysis used 1:1 nearest-neighbor matching without replacement. Both applied a caliper of 0.25 SD of the logit PS. Covariate balance is presented in fig. S8A, B.

Unadjusted ORs were estimated from matched 2×2 contingency tables. The Haldane-Anscombe correction (0.5 added to all four cells) was applied when any cell contained zero observations. Chi-squared tests were used for inference; Fisher’s exact test was substituted when any expected cell count was below five. P-values were FDR-adjusted (Benjamini-Hochberg) across all diseases within each comparison, with an adjusted threshold of 0.05. All analyses were conducted in R (version 4.4).

### Statistical analysis

Unless otherwise noted, all statistical tests were two-sided, and significance was defined as an FDR < 0.05. Multiple-testing correction was performed using the Benjamini–Hochberg procedure, applied within the relevant comparison group as specified in each subsection (e.g., within cohort × database for pathway enrichment, within comparison for EHR association testing). For within-versus between-tissue comparisons of NES profiles, one-sided Wilcoxon rank-sum tests were used, with effect sizes reported as median NES margins.

Differential gene expression significance was assessed using edgeR quasi-likelihood F-tests (disease signature generation) or DESeq2 (rat and mouse drug signatures), with genes considered significant at FDR < 0.05. Concordance between independently derived transcriptomic signatures (e.g., against DiSignAtlas) was quantified using Spearman correlation of gene-level log2 fold changes. Pathway-level enrichment was assessed via GSEA (fgsea), with pathways considered significant at adjusted p < 0.05; combination of enrichment evidence across signatures and definitions of tissue conservation, specificity, discordance, and LOO analysis are detailed under Gene Set Enrichment Analysis of Drug signatures. Reversal of disease signatures was quantified using a permutation-based Z-score of RGES derived from Kolmogorov–Smirnov connectivity statistics, with disease-level significance computed via adjusted Stouffer’s method (see Reversal of Disease-associated Signatures).

For EHR-based association analyses, unadjusted odds ratios were estimated from matched 2×2 contingency tables, with the Haldane–Anscombe correction (0.5 added to all cells) applied where any cell contained zero observations (*81*). Group differences were assessed using chi-squared tests, with Fisher’s exact test substituted when any expected cell count was below five. Missing covariate data (BMI) were imputed using multiple imputation by chained equations (MICE)(*80*).

All analyses were performed in R (v4.4), using the edgeR (v4), DESeq2, fgsea, msigdbr, STRINGdb (v12.0) packages.

## List of Supplementary Materials

Fig. S1 to S8

Tables S1 to S17

## Acknowledgments

We thank the constructive feedback from Drs. Charles Hong, Nagib Chalfoun, Jessica Devries, Reda Girgis, Marcia Gordon, and Dave Morgan. A.I. Use Disclosure: Claude (Anthropic) was used to assist with drafting and editing portions of the text and code. This work was supported in part through the Minerva computational and data resources and staff expertise provided by Scientific Computing and Data at the Icahn School of Medicine at Mount Sinai and supported by the Clinical and Translational Science Awards (CTSA) grant UL1TR004419 from the National Center for Advancing Translational Sciences

## Author contributions

Conceptualization: BC Work Coordination: BC

Disease Signature Database: SK, XZ EHR analysis: NT, BG

RNA-seq sample preparation: LH, XL Rat

Treatment: SW, AL, MM, HG

Bioinformatics support: RS, DJ, DL Writing: ZP, SK, NT, BC

## Data and materials availability

RNA-seq profiles of rat samples will be available on the Gene Expression Omnibus at GSE338416 once this paper is published. Other associated data generated during this study are included in the published article and the supplementary tables. Disease signature datasets available for query on the OCTAD 2.0 portal (https://apps.octad.org/octad2). The EHR data from Mount Sinai is not accessible to the public and can only be accessed by authorized individuals and hence cannot be shared directly.

Code is available on the OCTAD 2.0 github (https://github.com/Bin-Chen-Lab/octad2_GLP1).

