## Supplementary Figures for "In Vivo Transcriptomics-Based Framework for Systematic Indication Expansion"

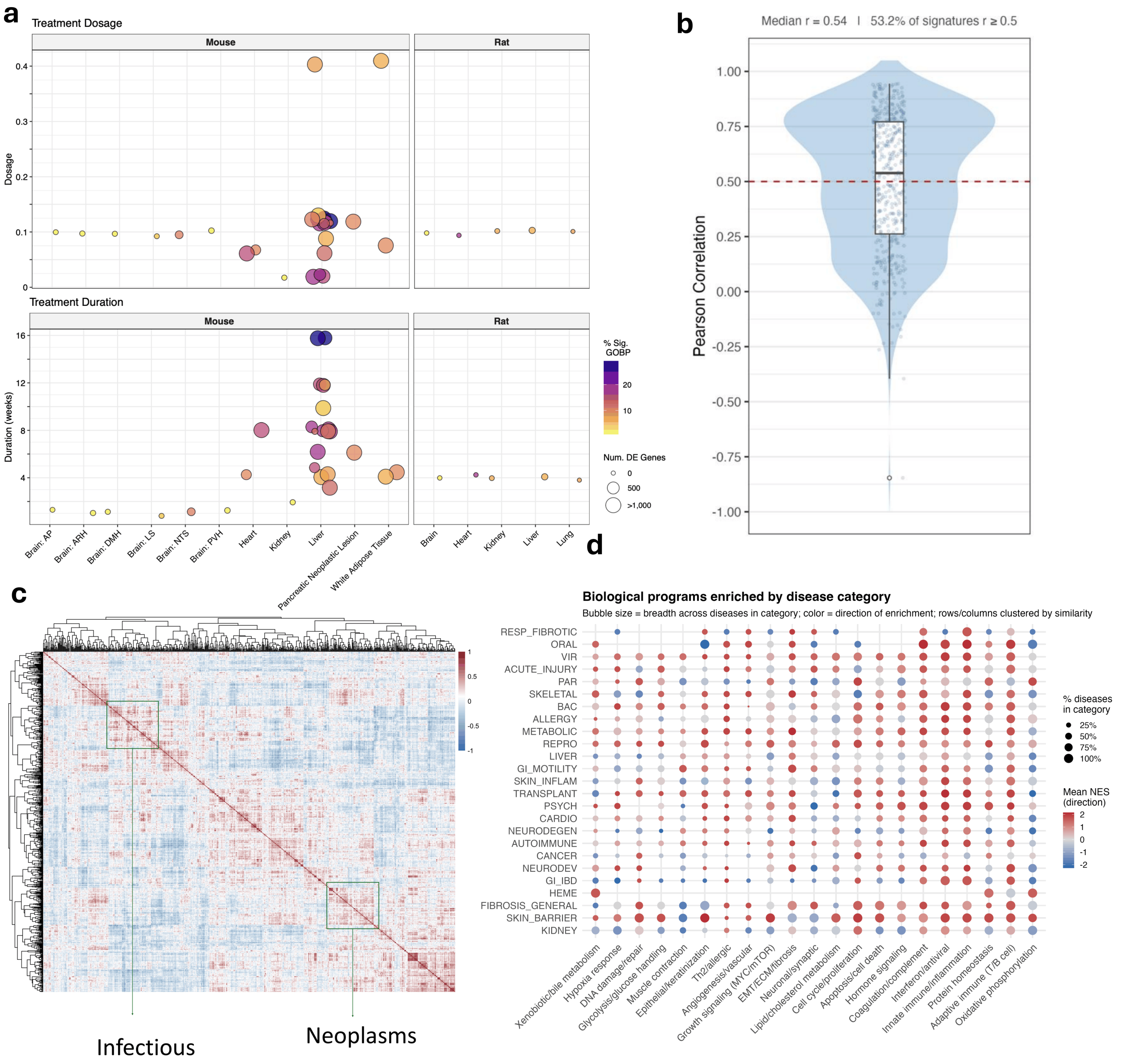


***fig. 1: Disease- and drug signatures.*** ***(A)*** *Bubble plots summarizing the 35 semaglutide-perturbed transcriptomic signatures across tissues and species (mouse and rat), ordered by treatment dosage and treatment duration. Each bubble represents a single signature; bubble size indicates the number of differentially expressed genes and bubble color indicates the percentage of significantly enriched Gene Ontology Biological Process terms (% Sig. GO:BP). Signatures with no detected differential expression are shown as minimal points. NA denotes signatures for which dosage or duration metadata were unavailable.* ***(B)*** *Violin plot showing the distribution of Pearson correlations between atlas disease signatures and matched disease signatures from the DiSignAtlas reference database. The dashed red line indicates r = 0.5; the median correlation and percentage of signatures exceeding this threshold are annotated. The concentration of*

*correlations above 0.5 demonstrates that atlas signatures are broadly concordant with independently curated reference signatures.* ***(C)*** *Pearson correlation across all diseases indicates clustering of similar disease types.* ***(D)*** *Concordance between enriched pathways and generally known disease mechanisms for disease groups, displayed as bubble plot. Columns are particular groups of pathways, rows are particular groups of disease signatures.*

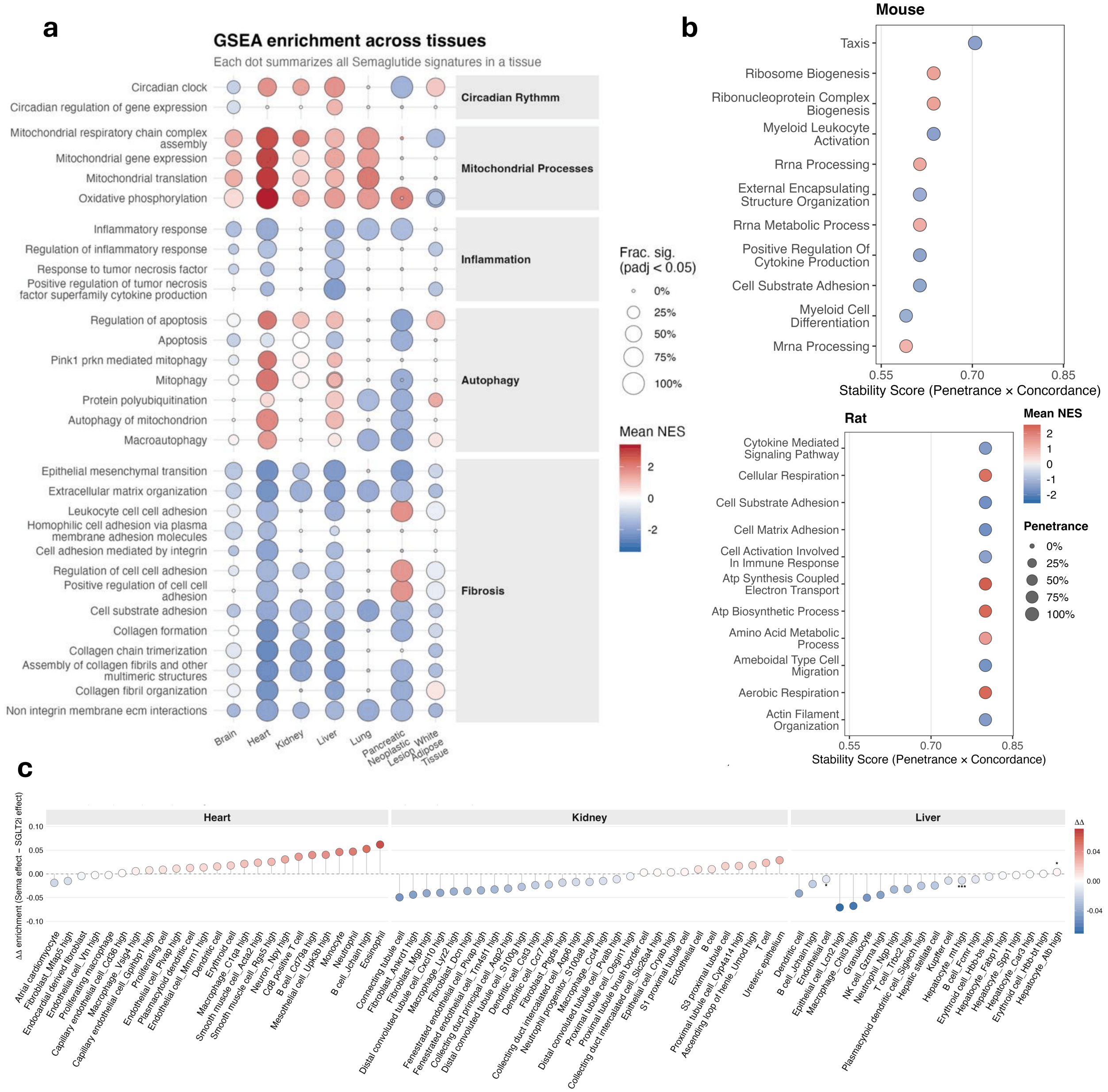


***fig. 2: Pathway and cell type changes associated with semaglutide treatment. (A)*** *Selected pathway enrichments of interest, including fibrotic processes, immune processes, mitochondrial and energetic processes, and circadian rhythm processes across tissues. Significant at the FDR < 0.05 level.* ***(B)*** *Lollipop plot of enrichments in the GO:BP database as a companion for the Reactome subfigure of Figure 2, across rats and mice.* ***(C)*** *A lollipop plot comparing cell type*

*populations changes in marker-based deconvolution of SGLT2 inhibitor and semaglutide-perturbed signatures. Points are equal to the mean of semaglutide minus the mean of SGLT2 inhibitor signature marker change across signatures; blue = lower under Sema, red = lower under SGLT2 inhibitor; asterisks indicate significance. The lack of differences between immune cells implies that in these tissues, semaglutide and SGLT2 inhibitors modulate immune cell populations similarly.*

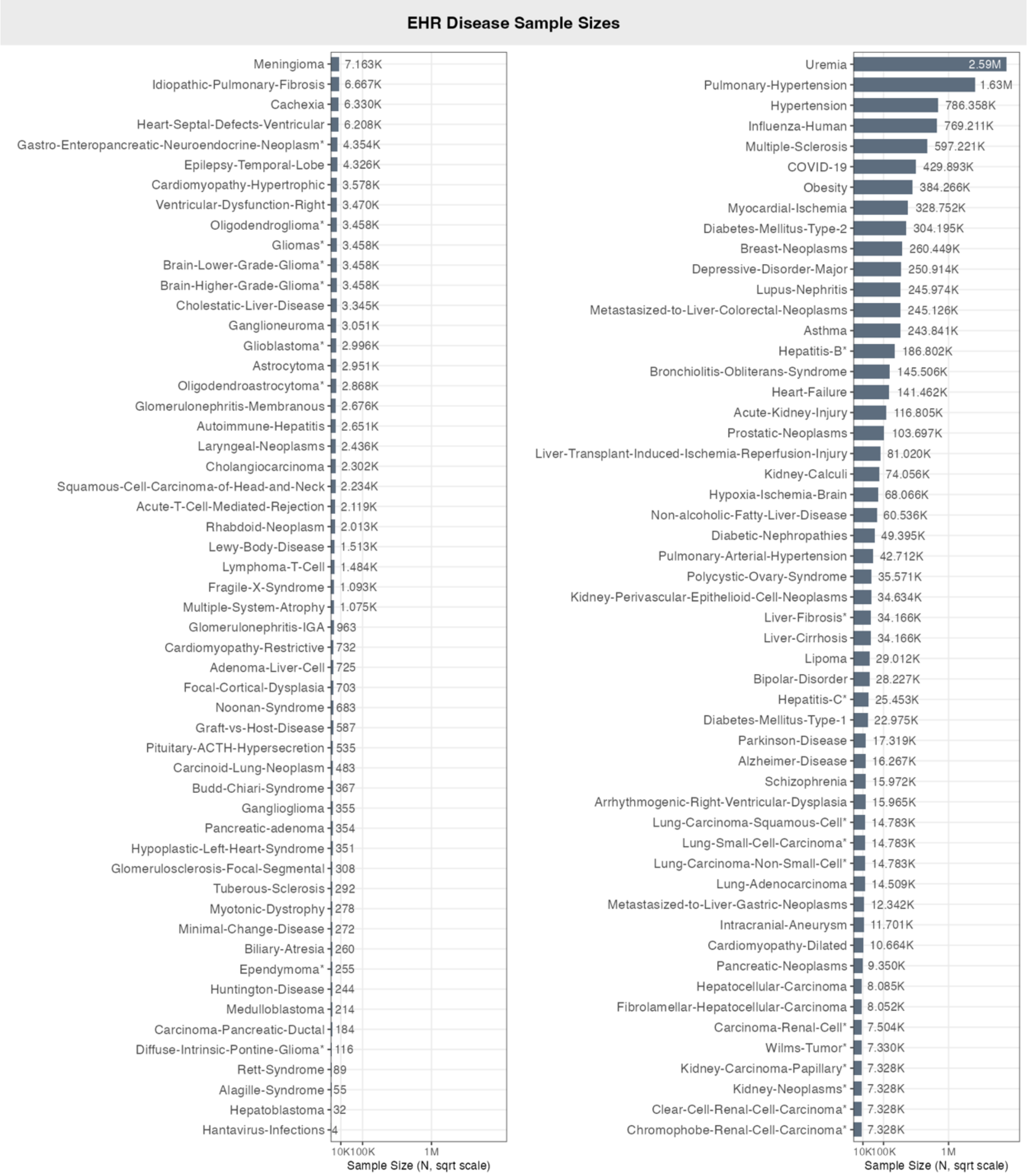


***fig. 3: Sample sizes of mapped conditions in Mount Sinai Health System.*** *Asterisks indicate multiple disease signatures mapped a single condition, see supplementary table 10 for exact mappings. The Mount Sinai Health System has records on 12,938,101 distinct patients as of July 29, 2026.*

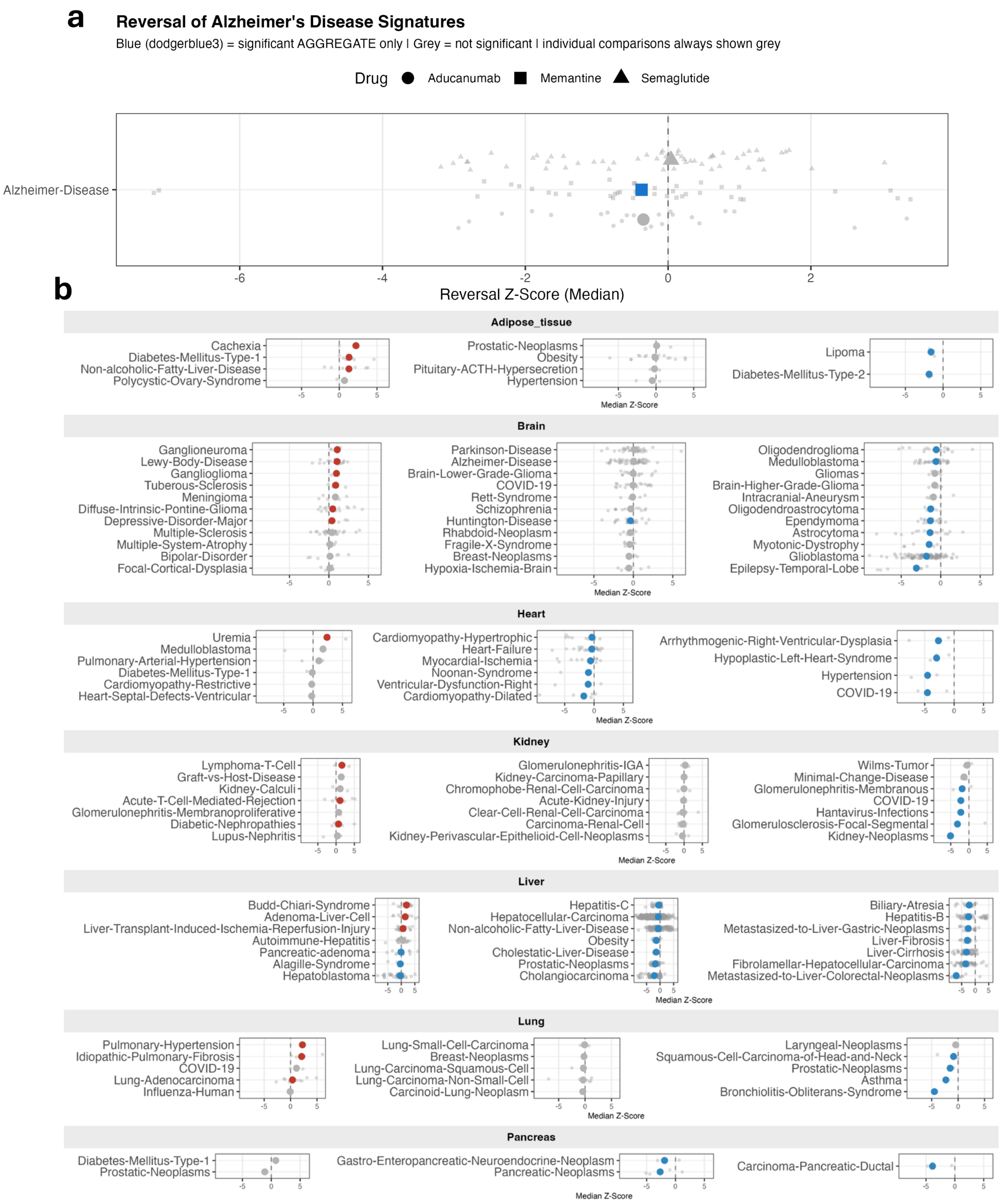


***fig. 4: Tissue-specific disease reversal by semaglutide and the comparison with other drugs in the reversal of Alzheimer's disease. (A)*** *Reversal of aducanumab and memantine signatures against Alzheimer's compared to semaglutide. Aducanumab's median Z-RGES score was numerically more reversed than semaglutide's (-0.346 vs. 0.042; FDR = 0.107 vs. 0.07), though this did not reach significance, likely owing to the relatively small current sample of isotype-control comparisons (n = 24) in comparison to memantine (n = 40). Despite aducanumab's lack of significance, the disease signatures most reversed by aducanumab had higher expression of the amyloid-beta clearance program (GO:BP; β = -2.39, p = 0.002), consistent with the known mechanism of promoting microglial clearance of Aβ, implying that aducanumab opposes, transcriptionally, the AD signatures showing the strongest evidence of active, unresolved*

*microglial clearance effort.* ***(B)*** *Reversal forest plots, visualized on a per-tissue basis, to contrast with the primary visualization in Fig. 3, in which reversal across tissue contexts is combined. It highlights the importance of the pan-tissue approach, as evidences from multiple sources combine into more accurate predictions. For instance, obesity is not strongly reversed in adipose tissue alone, possibly owing to a relatively smaller pool of comparisons for that tissue, but the addition of liver contexts ensures that on a systemic level the prediction reflects accurately.*

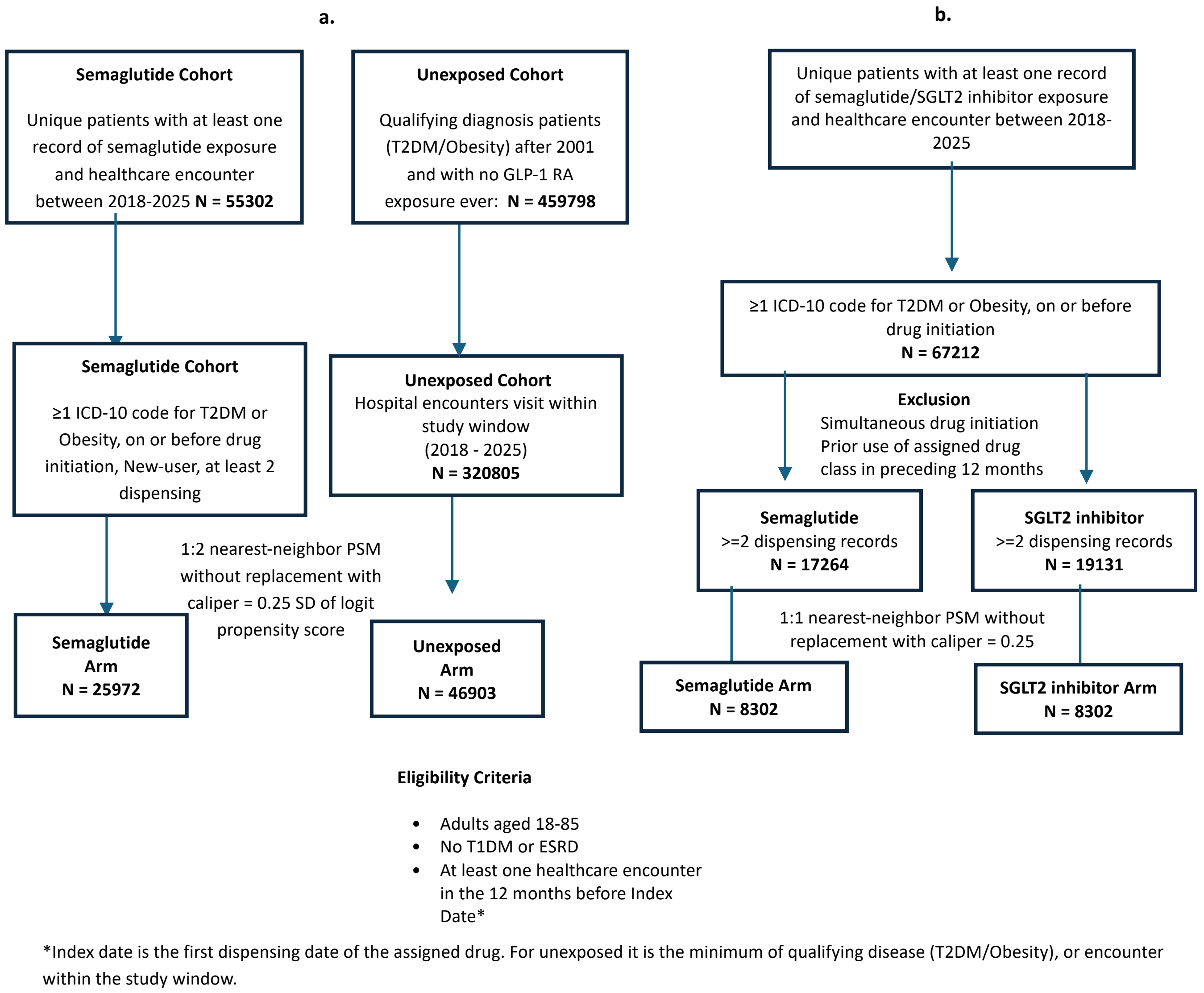


***fig. 5: Study design EHR analysis. (A)*** *Study design and patient selection workflow for the semaglutide vs unexposed patient groups analysis.* ***(B)*** *The corresponding study design and patient selection workflow for semaglutide vs SGLT2 inhibitor patient analysis.*

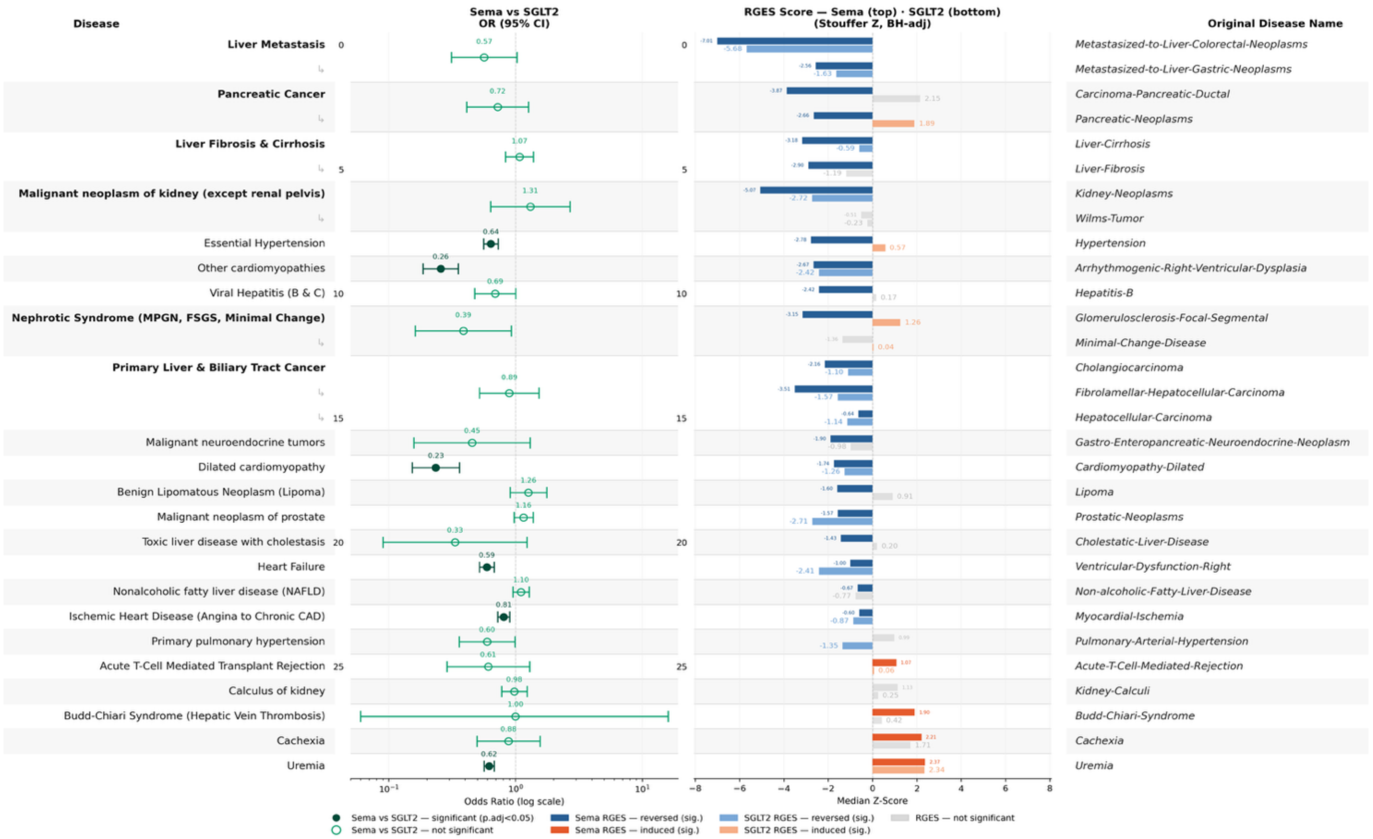


***fig. 6: Semaglutide vs SGLT2 inhibitors, comparison of reversal and EHR unadjusted odds ratios.*** *Real-world EHR data provides a valuable anchor, and this was the motivation for screening top reversal hits in the Mount Sinai Health System. Several significant associations were directionally concordant with RGES predictions in the semaglutide-unexposed comparison. Across the semaglutide-SGLT2 inhibitor comparison, all seven significant associations focused on cardiovascular and renal outcomes, reflective of the common indications shared between these drugs.*

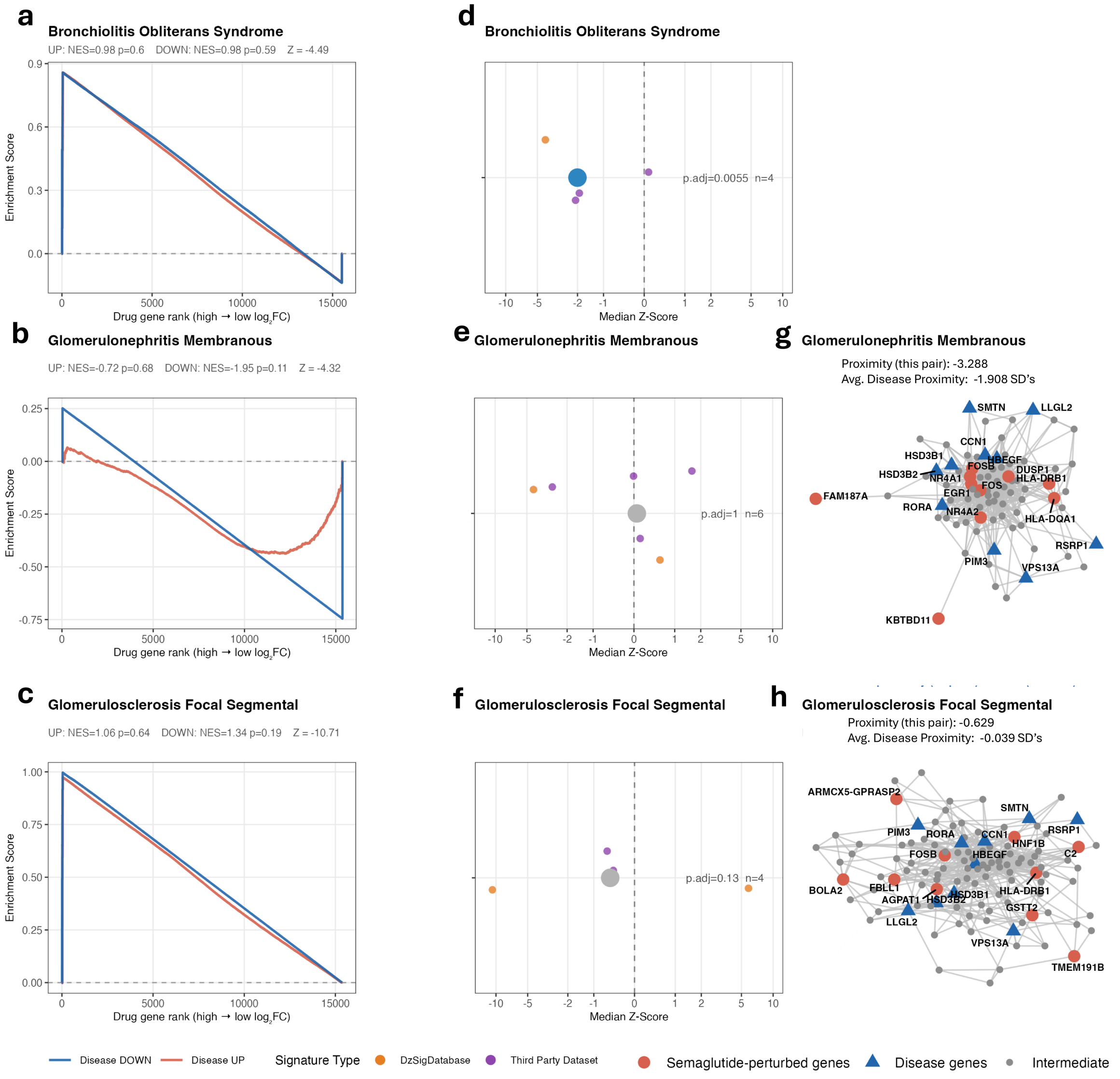


***fig. 7: Supplementary rare disease exploration. (A, B, C)*** *GSEA-style enrichment plots showing transcriptomic reversal of disease gene expression signatures by semaglutide for bronchiolitis obliterans syndrome (A) glomerulonephritis membranous (B) and glomerulosclerosis focal segmental (C). The blue curve traces enrichment of disease downregulated genes and the red curve traces enrichment of disease upregulated genes across the semaglutide-perturbed gene ranking (high to low log2FC). Crossing of the curves toward opposing ends of the ranked list indicates transcriptomic reversal. For bronchiolitis obliterans syndrome, the leading edge genes most responsible for reversal included genes of interest such as PCSK9, a major lipid-metabolism gene central to LDL cholesterol clearance, which has been noted to affect lung fibrosis, and NLRP2, an inflammasome-family gene, indicating possible modulation of lipid/immune function in the lung by semaglutide.* ***(D, E, F)*** *Distribution of Z-RGES scores across tissue-matched drug-disease comparisons for bronchiolitis obliterans (D)*

*glomerulonephritis membranous (E) and glomerulosclerosis focal segmental (F). Orange points represent comparisons from the harmonized disease atlas; purple points represent independent third-party disease signatures. The value of additional signatures becomes clear, as more contexts eventually demonstrated a lack of reversal for some diseases. Bronchiolitis obliterans syndrome is the exception of these three, although its signatures were derived from pseudobulk signatures from single-cell analysis, and are divided by GVHD (graft-vs host disease)-presenting bronchiolitis obliterans syndrome and bronchiolitis obliterans syndrome, across immune/non-immune cell fractions. The reversal was consistent across both fractions.* ***(G, H)*** *Protein-protein interaction network proximity plots for Glomerulonephritis Membranous (G) and Glomerulosclerosis Focal Segmental (H) derived from the most informative tissue-matched semaglutide signature for each disease. Red circles represent semaglutide-perturbed genes, blue triangles represent disease-associated genes, and grey nodes represent intermediate network connections. The proximity score indicates the mean shortest path distance between drug and disease gene sets in the network; values in parentheses indicate deviation from a degree-preserved random expectation in standard deviations, with more negative values indicating greater-than-expected network proximity. Bronchiolitis obliterans syndrome was not eligible for this adapted network analysis as the semaglutide perturbed lung signatures had no DE genes.*

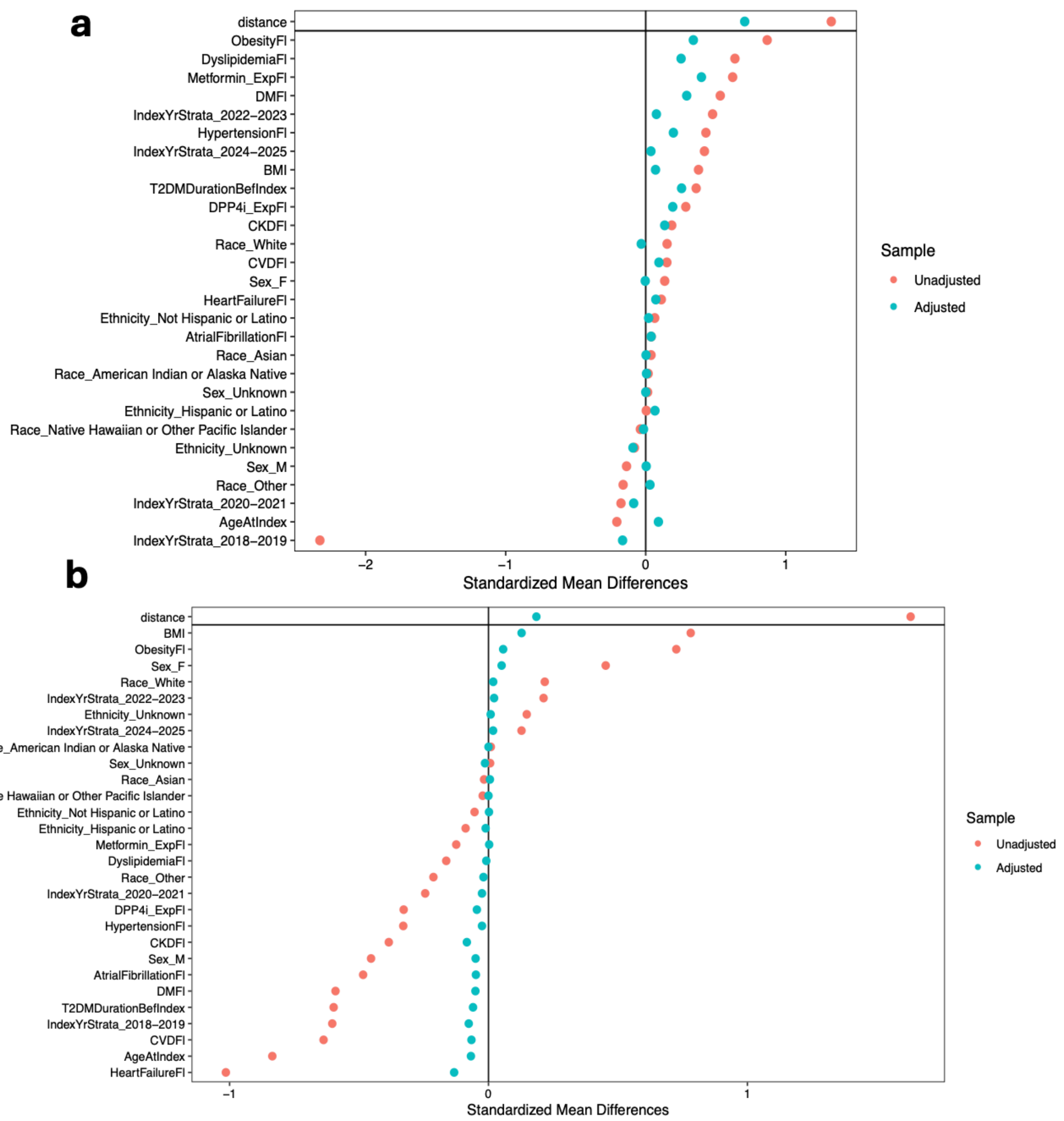


***fig. 8: Covariate balance for the semaglutide comparisons. a,** Love plot showing standardized mean differences for each covariate before (unadjusted) and after (adjusted) propensity-score matching of semaglutide-exposed vs. unexposed patients. **b,** Corresponding Love plot for semaglutide vs. SGLT2 inhibitor matching.*
